# Visible implant elastomer (VIE) success in early larval stages of a tropical amphibian species

**DOI:** 10.1101/2020.04.29.057232

**Authors:** Chloe Fouilloux, Guillermo Garcia-Costoya, Bibiana Rojas

**Affiliations:** Department of Biology and Environmental Science, University of Jyväskylä, Finland

**Author notes:** Corresponding Author: Chloe Fouilloux.

## Abstract

Animals are often difficult to distinguish at an individual level, but being able to identify individuals can be crucial in ecological or behavioral studies. In response to this challenge, biologists have developed a range of marking (tattoos, brands, toe-clips) and tagging (PIT, VIA, VIE) methods to identify individuals and cohorts. Animals with complex life cycles are notoriously hard to mark because of the distortion or loss of the tag across metamorphosis. In frogs, few studies have attempted larval tagging and none have been conducted on a tropical species. Here, we present the first successful account of VIE tagging in early larval stages (Gosner stage 25) of the dyeing poison frog (*Dendrobates tinctorius*) coupled with a novel anaesthetic (2-PHE) application for tadpoles that does not require buffering. Mean weight of individuals at time of tagging was 0.12g, which is the smallest and developmentally youngest anuran larvae tagged to date. We report 81% tag detection over the first month of development, as well as the persistence of tags across metamorphosis in this species. Cumulative tag retention versus tag observation differed by approximately 15% across larval development demonstrating that “lost” tags can be found later in development. Tagging had no effect on tadpole growth rate or survival. Successful application of VIE tags on *D. tinctorius* tadpoles introduces a new method that can be applied to better understand early life development and dispersal in various tropical species.

## Introduction

Distinguishing individuals within a population is often key in deciphering animal behavior. Animal identification has applications in understanding parental care (Ménard et al., 2001), migration dynamics (Matthews et al., 2011) and social hierarchies (Holekamp et al., 1997). Studies across the animal kingdom have developed methods that vary both in invasiveness and success (guppies: Croft et al., 2003; dolphins: Defan et al., 1990; bears: Diefenbach & Alt, 1998; salamanders: Osbourn et al., 2011) to allow researchers to differentiate between individuals within groups. If visual differentiation is not an obvious option, physical manipulation (e.g. toe clips, tattoos; Perret & Joly, 2002; Phillott et al., 2007) and tagging (i.e. passive integrated transponder (PIT), visible implant alphanumeric (VIA), visible implant elastomer (VIE); Phillott et al., 2007; Donnelly et al., 1994) have been the most commonly used methods implemented in mark-recapture studies.

Differentiating individuals is important when there is a lot of intrapopulation variation in behavior, and is becoming especially relevant as we begin to see individuals adapt to new challenges onset by the effects of global warming, habitat fragmentation, and human interactions. However, long-term marker-based studies have not often been applied to animals with complex life cycles, as the physical transformations induced with metamorphosis and growth generally entail the loss or distortion of the mark. In anurans, there have been a range of successful tagging methods both in adult and larval stages, but the diversity of larval tagging studies has been limited to common temperate species (e.g., Campbell Grant, 2008; Courtois et al., 2013), and very few studies have been able to create a methodology that spans the animal’s entire life cycle (Bailey, 2004; Bainbridge et al., 2015; Caballero-Gini et al., 2019; Campbell Grant, 2008; McHarry et al., 2018). The current reported success rate of tags could be inaccurate for species outside of temperate regions. Tadpoles, especially those from neotropical regions, are known to be tremendously plastic with respect to morphology (Touchon & Warkentin, 2008), time-to-metamorphosis (Rudolf & Rödel, 2007), and even pigmentation (McIntyre et al., 2004), all of which could affect tag success.

Understanding the dispersion dynamics and survival of anurans from aquatic to terrestrial habitats makes developmentally early larval tagging especially interesting. For example, many species of poison frogs have parental care where recently hatched tadpoles are transported from terrestrial sites to arboreal pools (Pašukonis et al., 2019; Ringler et al., 2013; Schulte & Mayer, 2017; Summers & Tumulty, 2013). Tadpole tagging could provide a quick and reliable method of following individuals across development, understanding relatedness within pools, and observing tadpole behavior and parental care in the field. Here, we use a novel anesthetic procedure for tadpoles followed by a VIE tag application on dyeing poison frog (*Dendrobates tinctorius*) larvae that is then monitored throughout development.

Larval anuran tagging has been limited with respect to both developmental stage and weight. Most of the larval tagging to date has been done beyond the point of the onset of hind leg development (Andis, 2018(Gosner stage 30); Bainbridge et al., 2015 (Gosner stage 36-38)). At this point *D. tinctorius* tadpoles are typically at least a month old, meaning they have already been transported by their fathers and have long since been subject to both predation risk and aggression by conspecifics (CF personal observation; Rojas, 2014, 2015; Rojas & Pašukonis, 2019). Therefore, in order to obtain more valuable life history information, tags need to be applied earlier in development.

To our knowledge, the developmentally earliest tagging study applied VIA/VIE tags around Gosner stage 25 (Courtois et al., 2013; Gosner 1960), but its application was limited to large temperate tadpoles (average weight around 1.5g) that could be manipulated in the field without anesthesia. In this study we use 2-phenoxyethanol, an anesthetic that does not need to be buffered and can be stored at room temperature (Acme-Hardestry, 2013; National Center for Biotechnology, 2020), making it field appropriate, and applied VIE tags to the smallest and developmentally earliest stages of larval anurans recorded to date. We follow growth rate and tag success across development, and discuss potential field applications in order to better understand the dispersion dynamics and behavior of protected frogs.

## Materials & Methods

### Study organism

We used tadpoles from a breeding laboratory population of *Dendrobates tinctorius* kept at the University of Jyväskylä, Finland. Adult pairs are each housed in a 55L terrarium that contains layered gravel, leaf-litter, moss substrate and is equipped with a shelter, ramps, and live plants. Terraria are maintained at 26C (±2C) and are automatically misted with reverse osmosis water four times a day, maintaining a humidity around 95% and lit with a 12:12 photoperiod. Frogs are fed live *Drosophila* fruit flies coated in vitamin supplements three times per week. Tadpoles are raised singly in 10 x 6.5 x 5 cm cups which are filled with spring water, and fed *ad libitum* a diet of fish food (JBL NovoVert flakes) three times a week. Adult and tadpole health and water levels are checked daily, and experimental tadpoles were weighed and photographed weekly.

### Anesthesia

Prior to tagging, tadpoles were anesthetized in a 14mL solution of a 1µl:1mL ratio of 2-phenoxyethanol (2-PHE) to spring water. 2-phenoxyethanol is an oily liquid at room temperature and does not need to be buffered for anesthetic purposes (Coyle et al., 2004). The solution was reused multiple times for multiple tadpoles within a single day of tagging; its effect did not deteriorate after multiple uses. Each day of tagging a new solution was made. Tadpoles were placed in anesthetic solution until there was no muscular contraction in response to agitation. This took approximately 3 minutes. The effect of anesthesia on tadpoles lasted approximately 6 minutes; within 10 minutes individuals had regained full muscular function. The effects of anesthesia were similar across developmental stages. We had no deaths in response to our anesthesia procedure, which was applied to a total of 40 individuals across both our pilot study and experimental manipulations; anesthetized tadpoles ranged across early larval developmental stages (Gosner 24-26).

### Tadpole tagging

We applied VIE tags to early larval stages of *D. tinctorius* and monitored tadpoles across development (Fig 2) to ensure the presence of the tags over time, and to test the effects of larval tagging and tag retention across metamorphosis. Previous studies reporting tadpole tagging have been done primarily with late-term tadpoles (Gosner stage 30+) whose snout-vent lengths (SVL) were double or triple the SVL of tadpoles in our experiment (Andis, 2018; Bainbridge et al., 2015; McHarry et al., 2018). Other studies also worked with amphibians who produce large egg clutches (*Litoria aurea*, 37000 eggs/clutch (Pyke & White, 2001); American bullfrog, 12000 eggs/clutch (Howard, 1978); *Alytes obstetricans*, 50 eggs/clutch (Reading & Clarke, 1988)), which allowed for large tag sample sizes (n = 53-90, depending on study). *Dendrobates tinctorius* lay clutches that range from 2 to 5 eggs with a high level of mortality (Rojas & Pašukonis, 2019). Due to the reproductive limitations of the system, our sample total (n = 27 tagged, n = 11 control) is less than previously published data.

Elastomer was mixed and loaded into syringes prior to each tagging session, according to the Northwest Marine Technology VIE tag protocol. Elastomer was stored in a freezer (−20C) during extended periods of disuse and in a refrigerator between individual tagging sessions; we found that mixed elastomer was no longer effective after a storage period of longer than three months. Average tagging procedure was executed in under 90 seconds. Throughout our pilot study we found that tag retention was most effective when placed dorsally; thus, this experiment only contained dorsally marked tadpoles. Each tadpole was marked only once.

Tadpoles were placed on a laminated surface and dried with a paper towel to improve grip; a needle was placed subcutaneously and dye (approx. 1 µl) was injected. For this experiment, we used a fluorescent green elastomer, though any color tag would have been suitable for application. Tadpoles were placed under UV light to ensure proper placement of the tag. Tadpoles post-anesthesia were placed in a pool of spring water and observed for 10 minutes to ensure proper return of muscular function. After the observation period, tadpoles were returned to the pool of water in which they were living.

### Statistical analysis

#### Tag retention and observation model

VIE tag retention and observation was modeled using a Bayesian CJS (Cormack-Jolly-Seber) survival model (see R and JAGS code in supplementary materials; Jolly, 1965; Lebreton et al., 1992; Seber, 1965). For each individual, we considered tag observation as a categorical variable that was recorded as absent (0) or present (1); tags that had been lost and not re-observed were marked NA after the last confirmed observation. We assessed the status of the tag and tadpole development (size, weight) weekly. Tag retention was recorded as present (1) for all weeks previous to the last observation and recorded as NA for all those that followed. Our coding schematic (see Fig 1) takes into account observer error as a tag that is not observed at one time point but seen later in development is recorded as “retained” throughout the entire unobserved period. The retention status of the tag is unknown after the last positive observation. Our model considered weekly discrete time steps where the retention and observation of the tag were considered latent variables that occurred with a certain probability (*ϕ* and *p*, respectively following the nomenclature commonly used for CJS models).

**Figure 1.**
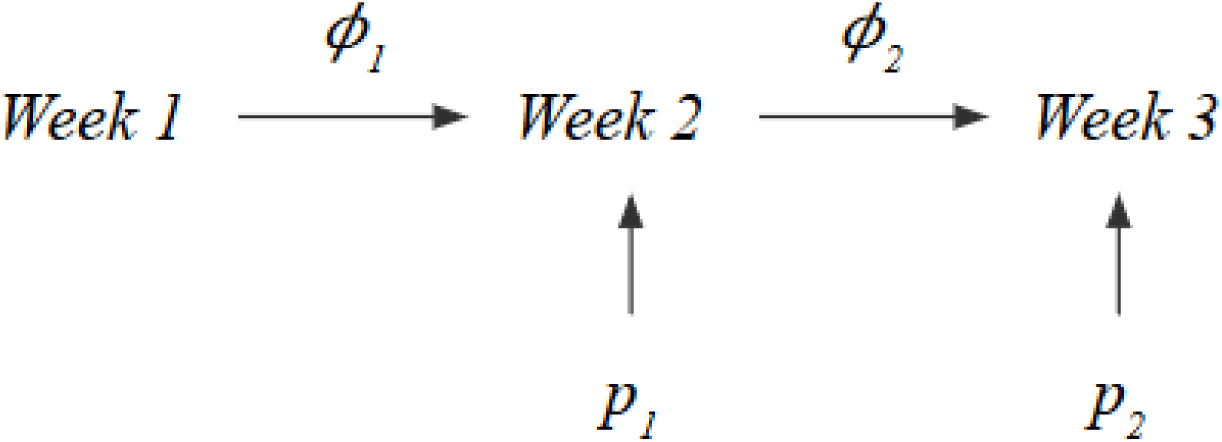
Schematic illustrating retention and observation variables in tag detection model. Where *ϕ* is the probability and *p* is the probability of observation. Consecutive weeks of an unobserved tag were marked “retained” if the tag was found later in development.

**Figure 2.**
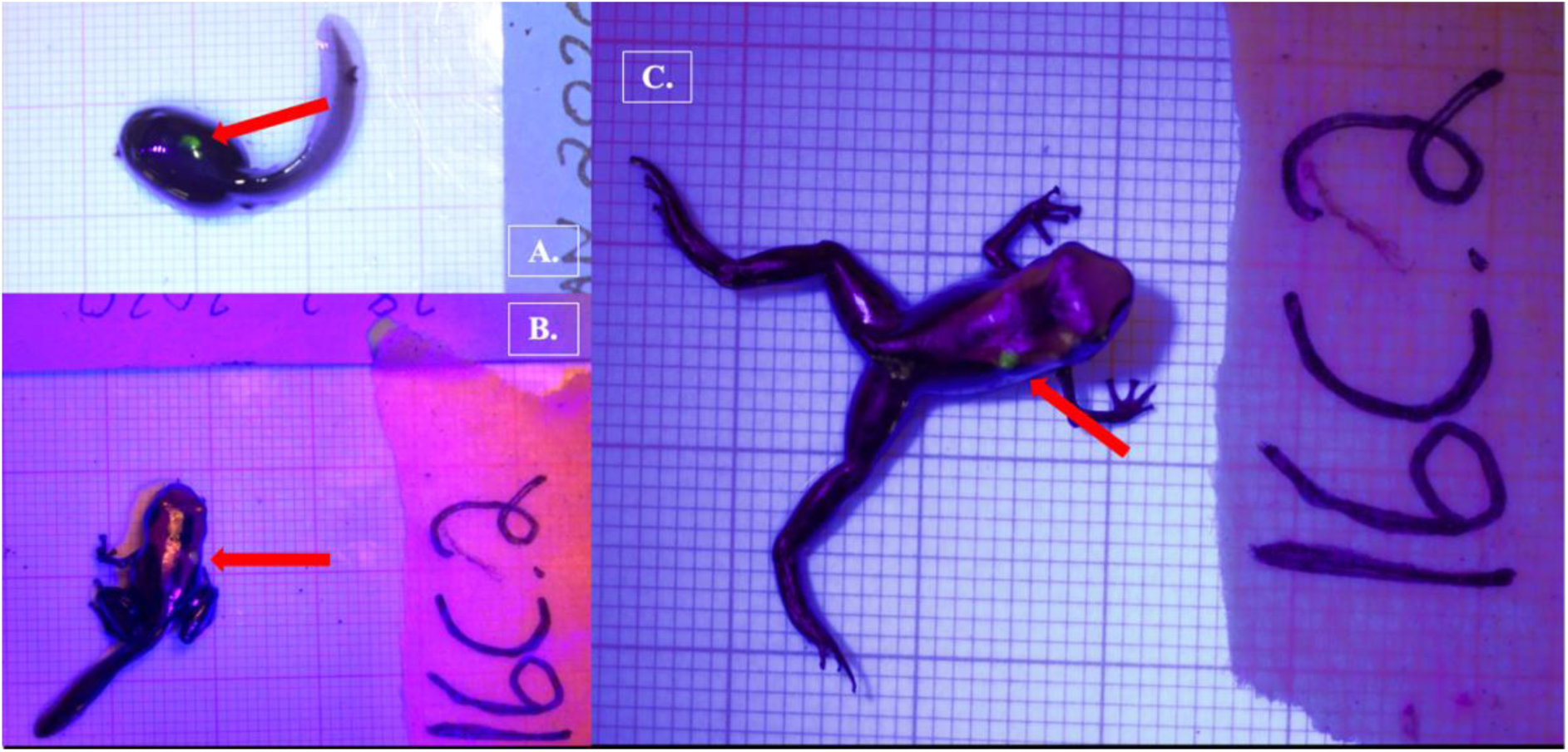
Fluorescent green VI Elastomer tag inserted dorsally on *Dendrobates tinctorius* shown on the same individual as (a) a late stage larva, (b) a metamorph, and (c) a recently metamorphosed juvenile. All photos taken with Nikon DS5300 DSLR on 1 x 1 mm background under UV light to enhance tag detection.

We considered three possible models (M1-3) of increasing complexity for *ϕ* and *p*: M1 assumed constant probabilities of observation and retention, M2 considered a week effect on both probabilities using a logit link function, and M3 took into account both a week effect on *ϕ* and *p* as well as individual identity as a random effect. Tag retention and observation were defined as following a binomial distribution with a probability *ϕ* and *p* respectively for all models. In models M2 and M3 retention and observation varied for each week of development (*t*), thus we used a logit link function to determine *ϕ* and *p* for each week considered. In model M3 retention and observation were also influenced by individual identity (*id)*, to account for it, we sampled the random effect parameter estimates from a normal distribution with a certain standard deviation for each individual which were later incorporated to the same logit link function.

For each model we used an MCMC approach considering uninformative priors for all parameters (see supplementary materials) and simulation run characteristics of 4 chains, 100,000 iterations with a 5,000 burn-in and a thinning of 10. Chain convergence was assessed using a potential scale reduction factor (PSRF) of our parameter estimates which discarded any model run that resulted in a PSRF larger than 1.1 or smaller than 0.9. We checked sample independence by determining the effective sample size of each parameter. We did not consider any model run with less than 5000 independent samples for any parameter.

Model selection was based on the lowest DIC value (Deviance Information Criterion) and biological relevance. The most likely discrete probabilities of retention and observation for each week were based on the posterior distributions generated by our model, these values were used to visualize the cumulative probability of tag detection across larval development.

#### Tadpole growth rates

Growth rates were compared between treatments using a linear mixed-effect model (LMM). Weekly weight (∼weight) and treatment (∼treatment) were coded as additive predictors in the growth rate model. Tadpole ID was used as a random effect on the intercept. Growth between treatments was compared by calculating weekly rate changes across development for both treatments, and then rate percent was used in model analysis which were evaluated with a Kenward-Roger’s method ANOVA. Growth rate models were chosen as a result of Akaike Information Criterion output (AIC; Akaike, 1973).

#### Tadpole survival rates

We used a Kaplan-Meier survival curve to visualize treatment effect on tadpole survival. A Cox proportional hazard model was used to calculate the parameters and uncertainty of tagging on survival. Survival object was parameterized with respect to death and time in response to treatment (Surv(Week, Dead) ∼ Treatment). Survival was coded as a binomial response (alive (0), dead (1)).

All models and statistics were performed in the program R using base R (v. 3.6.1, R Development Core Team, 2019) with additional packages “survival” (Therneau, 2014), “dplyr” (Wickham et al., 2019), “lme4” (Bates et al., 2015), “pbkrtest” (Halekoh & Højsgaard, 2014), “JAGS” (Plummer, 2003), and “R2jags” (Su & Yajima, 2015).

## Results

### Tag success

Out of our 27 fluorescent tags, 81 % (22/27) were successfully detected in tadpoles over the first month of application. This decreased to a little over 50% (8/15) detection by the third month of application, which also marks the approximate time of tadpole metamorphosis. Mean weight at time of tagging was 0.12g (± 0.019 SE) for tagged tadpoles and 0.099g (± 0.015 SE) for control tadpoles. Control tadpole weights ranged from 0.0307 to 0.18g at initial weigh-in, tagged tadpole weights ranged from 0.0318 to 0.36g at time of tagging. The smallest successful tag was applied at 0.0318g, which was a tadpole who had recently hatched (approximately Gosner stage 25). Our experimental tadpoles were tagged in the early larval stages of development: the youngest successful tag was applied on recently hatched tadpoles who had yet to be transported by their fathers. We report here that tagging did not prevent transport behavior by father (n = 2), though tagging at this life stage is especially delicate and requires a practiced hand. We attempted embryonic tagging in pilot studies, but were not able to successfully inject the tag without permanently damaging the embryo.

The model that had the lowest DIC did not include a week effect or an individual random effect (M1). It is important to note that tag observation sometimes changed throughout development, and tags that were not observed one week sometimes were detectable later in development (see Fig 3). Instances where tags were not observed could be due to individual growth, resulting in a tag being obstructed by a physical structure (i.e. muscle, tissue) for a period of time. For example, the cumulative probability of tag retention (*ϕ*) until the third month of development was 0.61 (0.33 -0.80, 95% CI) while the cumulative probability of tag observation (*p*) was 0.38 (0.16-0.59, 95% CI). On average, the difference between cumulative retention and observation rates was about 15%.

**Figure 3.**
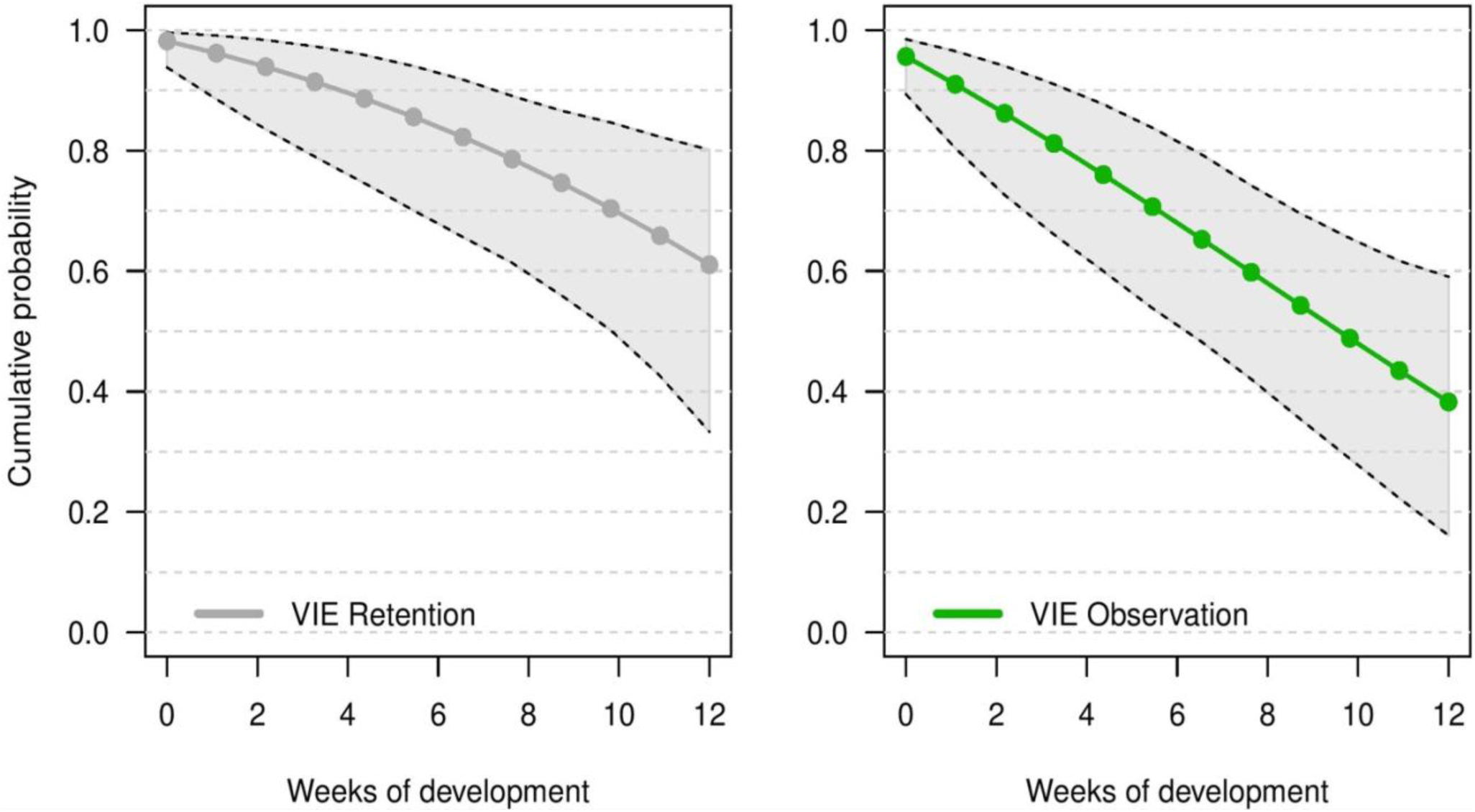
Estimate of the cumulative probability of tag detection across larval development for model M1. Points are posterior means. Grey points are the probability of tag retention (*ϕ*) and green points are probability of tag observation (*p*) at discrete time intervals corresponding to weeks of development. Grey polygons delimited by black dashed lines indicate the 95% credible intervals.

### Growth rate

We found no significant difference in weekly growth rate between control and tagged tadpoles (Fig 4), indicating that tagging does not affect tadpole growth (lmer, ANOVA Kenward-Roger’s method, F(1, 37) = 1.12, p = 0.296). Weekly tadpole growth rate significantly decreased across time (lmer, ANOVA Kenward-Roger’s method, F(1, 415) = 56.4, p = 0.03563^-11^).

**Figure 4.**
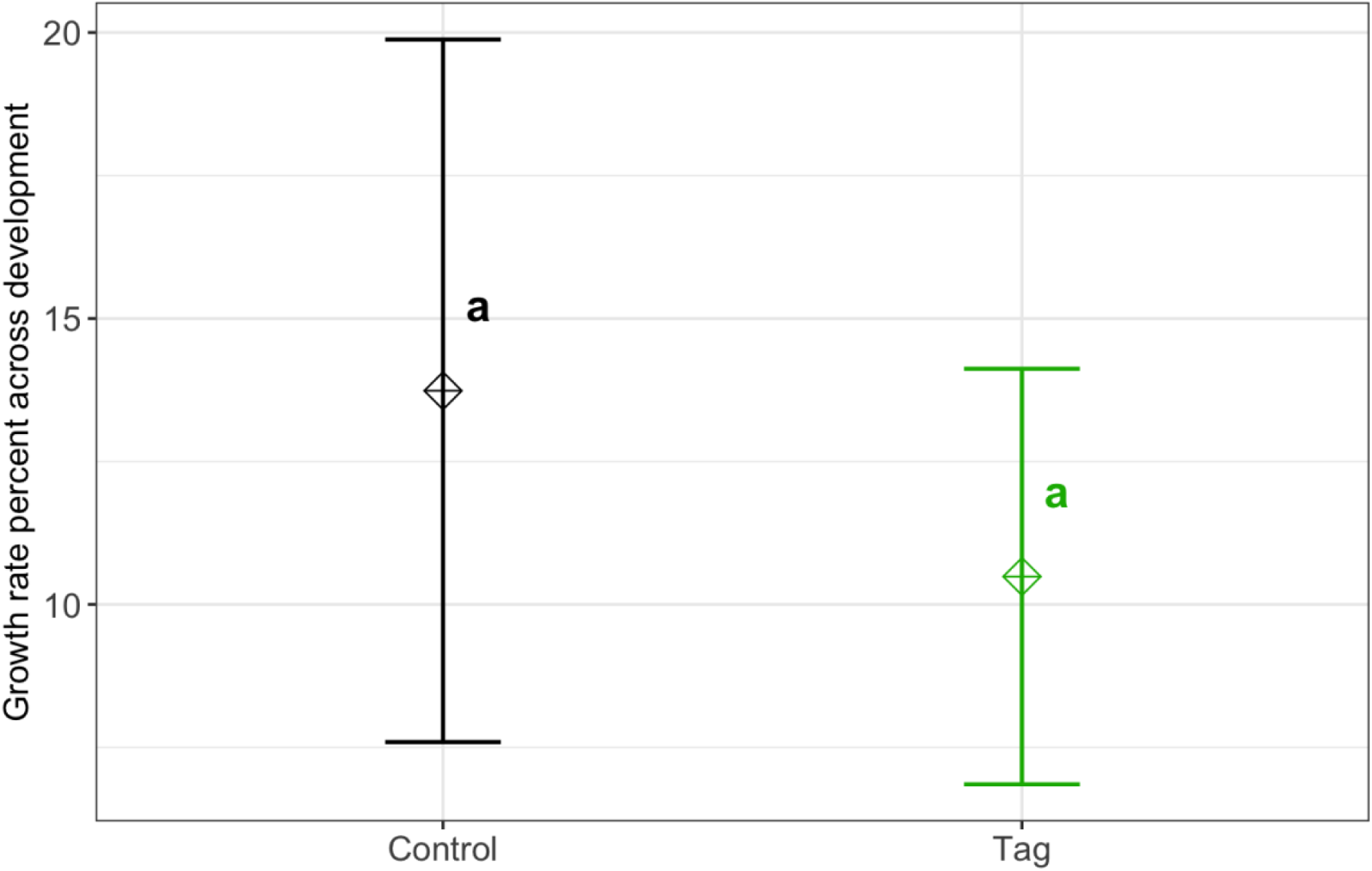
Average growth rate percent of VIE tagged and control groups. Diamonds represent the LS mean for which error bars indicate 95% confidence intervals. Means sharing letters are not significantly different (Tukey-adjusted comparisons, ANOVA Kenward-Roger’s method, F(1, 37) = 1.12, p = 0.296).

### Survival

There was no significant difference in survival between control and tagged groups across larval stages of development. Mortality across the first three months was 18% (n =5/27) for tagged tadpoles and 27% (n = 3/11) for control tadpoles (Fig 5). A Cox proportional hazard model did not find any significant difference in survival based on treatment (Cox proportional hazards model, z = -0.29, p = 0.76). Post-metamorphic survival was excluded from analysis due to unnaturally high froglet loss throughout the lab colony which is not indicative of tag impact on froglet survival, but likely ineffective laboratory practices for juvenile health.

**Figure 5.**
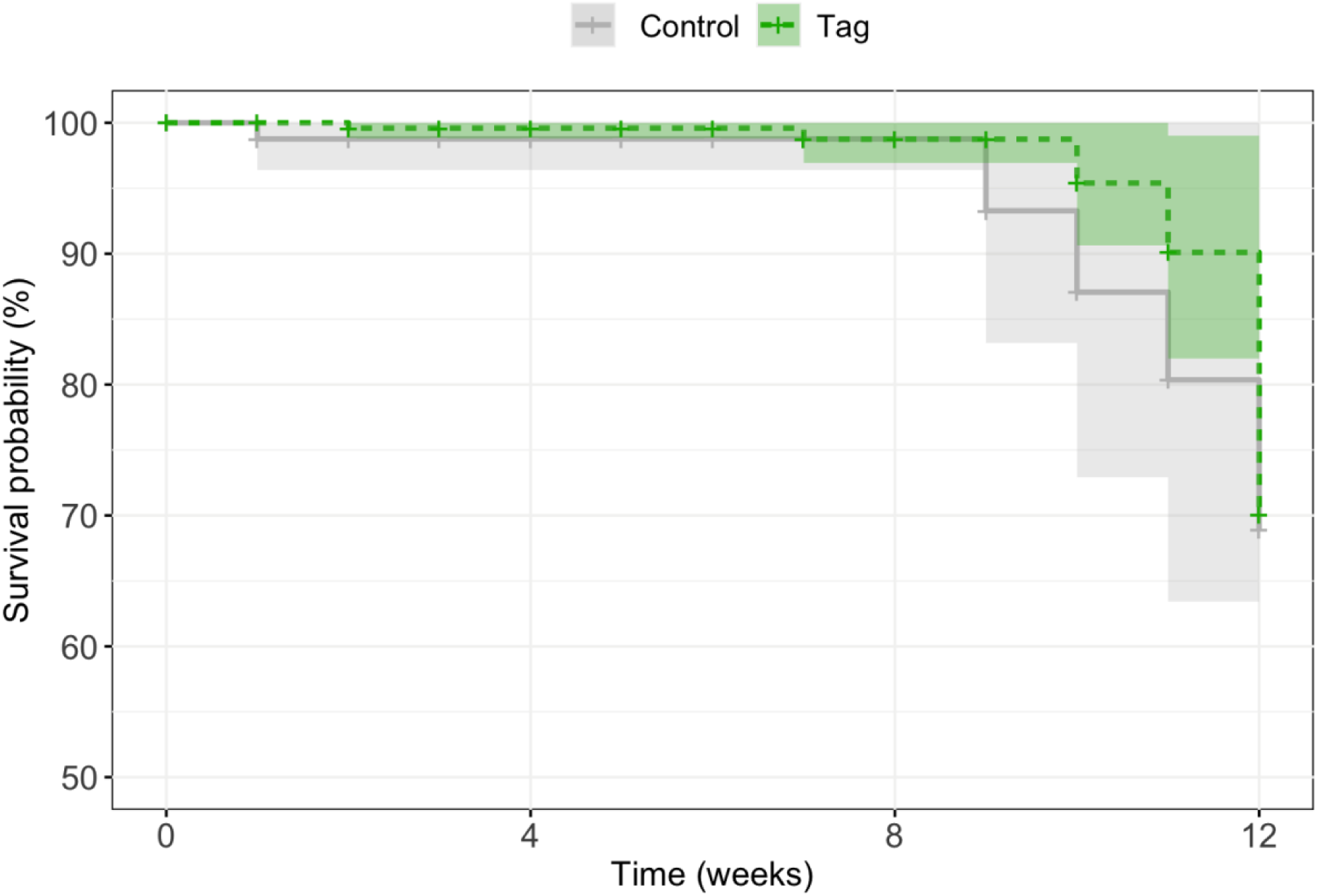
Kaplan Meier survival analysis of experimental tadpoles across larval development. A Cox proportional hazards model did not find any significant difference in treatment on tadpole survival across development (z = -0.29, p = 0.76).

## Discussion

In our study we applied VIE tags on *D. tinctorius* tadpoles and monitored them across development under laboratory conditions. Compared to previously published visible implant studies, our approach presents application at the youngest developmental stage, and is only the third study (after Bainbridge et al., 2015; Andis, 2018) to follow tags across metamorphosis. To our knowledge, this is the first attempt of larval tagging in a tropical frog, as previous work focused exclusively on species from temperate regions (Bainbridge et al., 2015: *Litoria aurea*; Courtois et al., 2013, *Alytes obstetricans;* Nauwelaerts et al., 2000, *Rana esculenta*). The successful application of VIE tags for the first time in a tropical species with elaborate parental care provides valuable opportunities to investigate parent-offspring interactions and dispersion of these species in a natural context.

Similar to other studies, we found no difference in growth rate or survival between tagged and control treatments. Based on our weekly weigh-ins and LMM model, we did not detect any significant impact of tagging on growth rates across development. Throughout our study, tadpoles grew significantly faster earlier in development which could be due to laboratory conditions (a high-food, no-competition environment). Given the circumstances, tadpoles may have invested energy in growing early in development which would help avoid predation and decrease latency to metamorphosis in the wild (Caldwell & De Araújo, 1998; Rojas, 2014). We found no effect of tagging on *D. tinctorius* survival across development; however, we had high rates of post-metamorphic mortality across our laboratory population. Natural history studies of *D. tinctorius* have shown high larval death rates (Rojas & Pašukonis, 2019). Although our tadpoles were not subject to the same pressures as wild populations, the mortality we observed across both treatments reflect the precariousness of early life stages in *D. tinctorius*.

After the first month of observation, 81% of tadpoles retained their tag, which is on par with retention rates reported in other tagging efforts (Anholt et al., 1998; Martin, 2011). Other studies report even higher rates of success with tadpoles elastomer tags (Courtois et al., 2013 (100%), Bainbridge et al., 2015 (100%)) which could be due to a shorter larval period and larger individuals at time of application. Retention rates of this study are of note because VIE tags have been extensively used for mark-recapture studies in fishes and anurans; when taking into account our tag retention rate and our tagging procedure (which takes less than 90 seconds), we can conclude that larval VIE mark-recapture studies on tropical amphibians is feasible.

A relevant note about implant tagging is that it is limited to observer perception. As tadpoles develop, morphological and phenotypic individual changes can facilitate or mask the presence of a tag. Most importantly, the lack of tag observation should not be assumed to indicate tag loss. In our experiment we were able to differentiate the cumulative probability of retention versus observation over time as a result of weekly checks of tag condition in experimental tadpoles. Over three months of development tags could go multiple weeks unobserved; finding them later in development indicated that tags were not lost, but had shifted position or been re-exposed as a result of growth. This is important to take into account for mark-recapture studies in settings where regular sampling or capture of the entire tagged population isn’t feasible. Our model estimates an average 15% difference in tag observation versus retention across larval development which is an error that can be incorporated as an informative prior in future tagging studies.

VIE tags come in a range of fluorescent colors, making the distinction of clutches or individuals from a distinct cohort possible. This is especially relevant for the larval stages of *D. tinctorius* when tadpoles are aggressive cannibals, as tagging efforts would help distinguish resident tadpoles in phytotelmata. Thus, tagging could be used to help monitor who is being deposited and who is getting attacked, allowing us to track interactions between tadpoles in ephemeral pools (Rojas, 2015). Moreover, VIE tagging of *D. tinctorius* makes it possible to successfully tag tadpoles before they are picked up and transported by their parent. Elastomer tags most clearly fluoresce under low-light conditions, making them ideal for their application in wild D. *tinctorius* tadpoles which live in dimly lit closed canopy rainforest.

VIE tags are one of the smallest tagging methods available for field studies. With respect to other tagging methods, VIA tags require a minimum SVL of 2 cm (Courtois et al., 2013) and PIT tags require 4 cm (Courtois et al., 2013), making VIE tags a unique option to study larval dynamics. VIE tags are not more than 4 mm in length, meaning that their successful application presents new opportunities to study larval amphibians that may not have been considered in the past. For example, *Anomaglossus beebei*, a small endemic poison frog from Guyana, has been seen to transport tadpoles multiple times throughout development (CF, personal observation). Larval tagging of this species could help decipher how shifting male territories influences larval care and transport, and if newly established males take care of tadpoles that are not their own. Early larval tagging could also work for *Allobates femoralis*, another tadpole transporter, to understand the shifting genetic diversity within phytotelmata across time (Erich et al., 2013).

Coupled with the unique patterning of *D. tincorius* that emerges in late metamorphosis and settles in adulthood (Courtois et al., 2012; Rojas & Endler, 2013), tags can provide early life identification that could be followed by pattern recognition, enabling individual discrimination throughout an individual’s entire lifespan. Bainbridge et al. (2015) report recently metamorphosed VIE tag retention to be high (88-95%); we also find that tags that lasted throughout larval development persisted across metamorphosis and into terrestrial life. Aside from implant tagging, genetic tracking has proven to be a reliable method to follow amphibian larvae throughout development into adulthood. With this said, genetic tracking does not provide immediate individual detection; further, studies using this method have been limited to individuals in a closed population, making the recapture of (surviving) tracked individuals reasonably certain (Ringler et al., 2015). In *D. tinctorius*, however, males can travel remarkable distances while carrying tadpoles (Pašukonis et al., 2019) making genetic tracking a less suitable method for individual distinction in this species. Andis (2018) also did remarkable work dyeing tadpoles of *Rana sylvatica* with calcein. This dye appears to persist across metamorphosis, though it should be noted that their development is much shorter than *D. tinctorius* and staining only allows for presence/absence detection. The presence of a VIE tag (and the range of colors available for application) allows for immediate discrimination of multiple groups/cohorts which may be an important advantage when conducting behavioral experiments and elucidating natural history dynamics in the wild.

Our study presents a successful continuation expanding marking methodology to larval tropical species. Using laboratory conditions, we were able to mimic a common scenario where experimental tadpoles were left to develop in small, stable pools of water. This is reflective of the most common parental behavior exhibited by *D. tinctorius*, where males transport newly-hatched tadpoles to develop in small water holdings (Rojas & Pašukonis, 2019). Future studies in field conditions would be useful to supplement these findings. For example, it will be important to understand how tadpole interaction with conspecifics, heterospecifics, and predators affects tag success. However, based on previously published data and the observation rates of our elastomers in this experiment, we believe that the application of elastomers in the wild is already an appropriate method to distinguish tadpoles for behavioral experiments. Elastomers are small, successful, and relatively easy to apply in early amphibian life stages. Our study contributes to the growing body of methods-based research demonstrating that visible implant elastomers are a viable tagging solution on a variety of anuran species in early development.

Ethics statement: This experiment was permitted by the National Animal Experiment Board (ESAVI/9114/04.10.07/2014). Raw data and R code will be available upon acceptance at the University of Jyväskylä data repository. DOI xxx.

## Conclusions

Differentiating individuals/cohorts can be a powerful tool when conducting behavioral experiments. Often, marking animals is a technique used to distinguish individuals when physical features are not distinct enough for visual differentiation. Choosing an optimal tag for a system is a tradeoff between reliability and invasiveness and is often limited to product cost and efficiency in identification. Elastomers (VIE) are injectable polymers that have been extensively used in fish and anuran systems. However, until this point, they have been applied to large larvae or adults and have been heavily biased towards common, temperate species. Here, we present the first application of VIE tags on a small larval tropical frog (*Dendrobates tinctorius*) and follow tag success across development. We found that (1) VIE tags can be successfully applied to recently hatched tadpoles, (2) tags can be reliably followed throughout larval development and sometimes retained across metamorphosis, and (3) VIE tags do not appear to interfere with parental care behavior (i.e. tadpole transport). Our study expands the application of tagging to early developmental stages in tropical amphibians which can be of use in behavior, conservation, and natural history research studies in the future.

## Supporting information

Raw Data

VIE Growth/Survival Code

VIE Model Code

## Acknowledgements

We would like to give an enormous thank you to Matthieu Bruneaux for his conversation and recommendation with respect to Bayesian modeling. We also owe a big *kiitos* to master’s students Nina Kumpulainen and Emmi Alanen and lab technician Teemu Tuomaala for helping with tadpole care--it takes a village.

## Notes

### Competing Interest Statement

The authors have declared no competing interest.

