## Supplementary Tables/Figures for "Visible implant elastomer (VIE) success in early larval stages of a tropical amphibian species"

*Supplementary Materials*

Table 1. Mathematical formulation for each of the models considered in JAGS.

| Model | Mathematical formulation |
| --- | --- |
| M1 | $Retention \sim B\left( \phi\right)$ |
|  | $Observation \sim B(p)$ |
| M2 | $\phi_{t}=\frac{1}{1+e^{-(\alpha_{\phi}+\beta_{\phi}t)}}$  ${Retention}_{t} \sim B\left( \phi_{t} \right)$ |
|  | $p_{t}=\frac{1}{1+e^{-(\alpha_{p}+\beta_{p}t)}}$  ${Observation}_{t} \sim B\left( p_{t} \right)$ |
| M3 | $\alpha_{\phi_{id}}\sim N(0,\sigma_{\alpha_{\phi_{id}}}^{2})$  $\beta_{\phi_{id}}\sim N(0,\sigma_{\beta_{\phi_{id}}}^{2})$  $\phi_{t,id}=\frac{1}{1+e^{-(\alpha_{\phi}+\alpha_{\phi_{id}}+\left( \beta_{\phi}+\beta_{\phi_{id}} \right) t)}}$  ${Retention}_{t,id} \sim B\left( \phi_{t,id} \right)$ |
|  | $\alpha_{p_{id}}\sim N(0,\sigma_{\alpha_{p_{id}}}^{2})$  $\beta_{p_{id}}\sim N(0,\sigma_{\beta_{p_{id}}}^{2})$  $p_{t,id}=\frac{1}{1+e^{-\left( \alpha_{p}+\alpha_{p_{id}}+\left( \beta_{p}+\beta_{p_{id}} \right) t \right)}}$  ${Observation}_{t,id} \sim B\left( p_{t,id} \right)$ |

Table 2. Posterior distribution results of the M1 model

| Parameter | Mean | SD | 2.5% Q. | 97.5% Q. |
| --- | --- | --- | --- | --- |
| $\phi$ | 0.9572 | 0.01301 | 0.9293 | 0.9800 |
| $p$ | 0.9274 | 0.01775 | 0.8904 | 0.9581 |
| Deviance | 118.3336 | 8.56818 | 105.6864 | 137.3522 |

Table 3. Posterior distribution results of the M2 model

| Parameter | Mean | SD | 2.5% Q. | 97.5% Q. | Significance |
| --- | --- | --- | --- | --- | --- |
| $\alpha_{\phi}$ | 3.08207 | 0.51868 | 2.127 | 4.1696 | * |
| $\beta_{\phi}$ | 3.99445 | 0.72535 | 2.7159 | 5.56701 | * |
| $\alpha_{p}$ | -0.09068 | 0.08951 | -0.2685 | 0.08852 |  |
| $\beta_{p}$ | -0.12104 | 0.11338 | -0.3400 | 0.10763 |  |
| Deviance | 120.53643 | 9.63689 | 105.7178 | 142.22163 |  |

Table 4. Uninformative priors used for every parameter & model. * Indicates that only positive values were sampled from the normal distribution for the random effects model. The standard deviation of the normal distributions from which the random effect of individual identity were sampled can only assume positive values.

| Model | Parameter | Prior distribution |
| --- | --- | --- |
| M1 | $\phi$ | Uniform distribution with 0 and 1 as limits |
|  | $p$ | Uniform distribution with 0 and 1 as limits |
| M2 | $\alpha_{\phi}$ | Normal distribution, mean = 0, sd = 10^-4^ |
|  | $\beta_{\phi}$ | Normal distribution, mean = 0, sd = 10^-4^ |
|  | $\alpha_{p}$ | Normal distribution, mean = 0, sd = 10^-4^ |
|  | $\beta_{p}$ | Normal distribution, mean = 0, sd = 10^-4^ |
| M3 | $\alpha_{\phi}$ | Normal distribution, mean = 0, sd = 10^-4^ |
|  | $\beta_{\phi}$ | Normal distribution, mean = 0, sd = 10^-4^ |
|  | $\alpha_{p}$ | Normal distribution, mean = 0, sd = 10^-4^ |
|  | $\beta_{p}$ | Normal distribution, mean = 0, sd = 10^-4^ |
|  | $\sigma_{\alpha_{\phi}}$ | Normal distribution*, mean = 0, sd = 10^-4^ |
|  | $\sigma_{\beta_{\phi}}$ | Normal distribution*, mean = 0, sd = 10^-4^ |
|  | $\sigma_{\alpha_{p}}$ | Normal distribution*, mean = 0, sd = 10^-4^ |
|  | $\sigma_{\beta_{p}}$ | Normal distribution*, mean = 0, sd = 10^-4^ |

Table 5. Chain convergence estimated through the Potential scale reduction factor (PSRF) and effective sample size of the parameters from models M1 and M2.

| Model | Parameter | PSRF (Point est.) | PSRF (Upper C.I.) | Effective Size |
| --- | --- | --- | --- | --- |
| M1 | $\phi$ | 1 | 1.00 | 2054.260 |
|  | $p$ | 1 | 1.00 | 1802.229 |
| M2 | $\alpha_{\phi}$ | 1 | 1.00 | 1276.573 |
|  | $\beta_{\phi}$ | 1 | 1.00 | 1145.789 |
|  | $\alpha_{p}$ | 1 | 1.01 | 1311.156 |
|  | $\beta_{p}$ | 1 | 1.01 | 1240.023 |

**A**
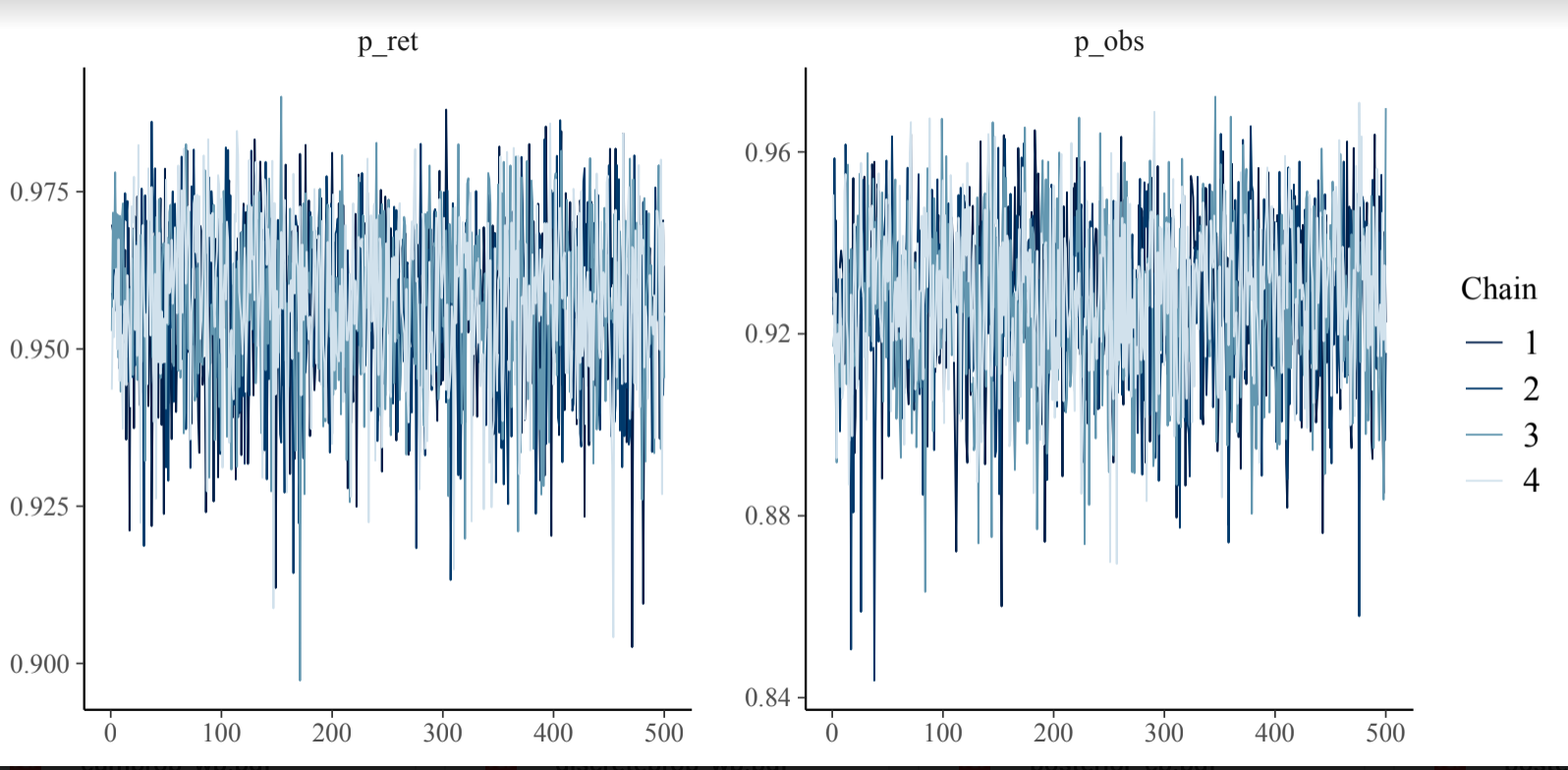
. **B**.
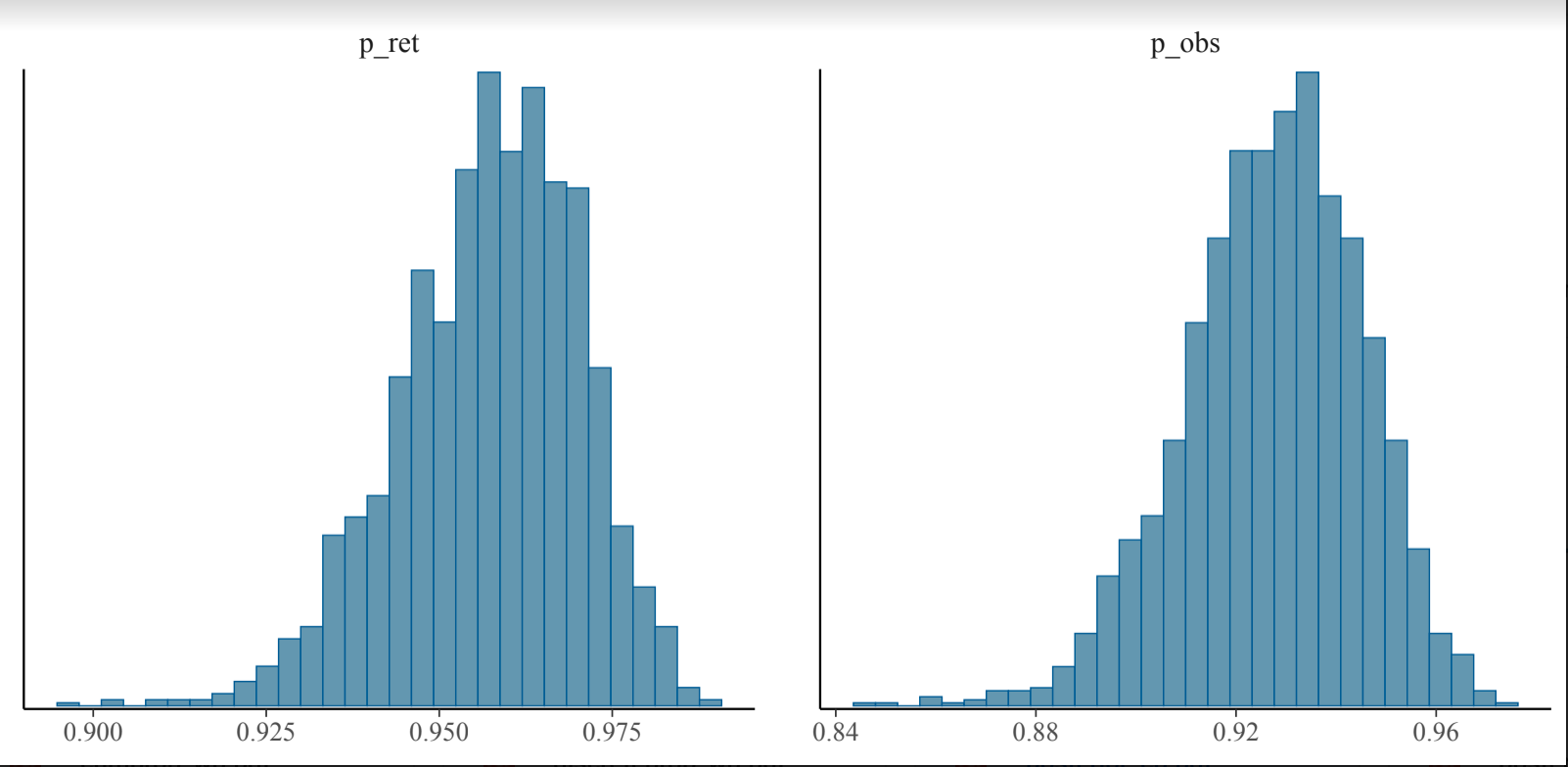
**C.**
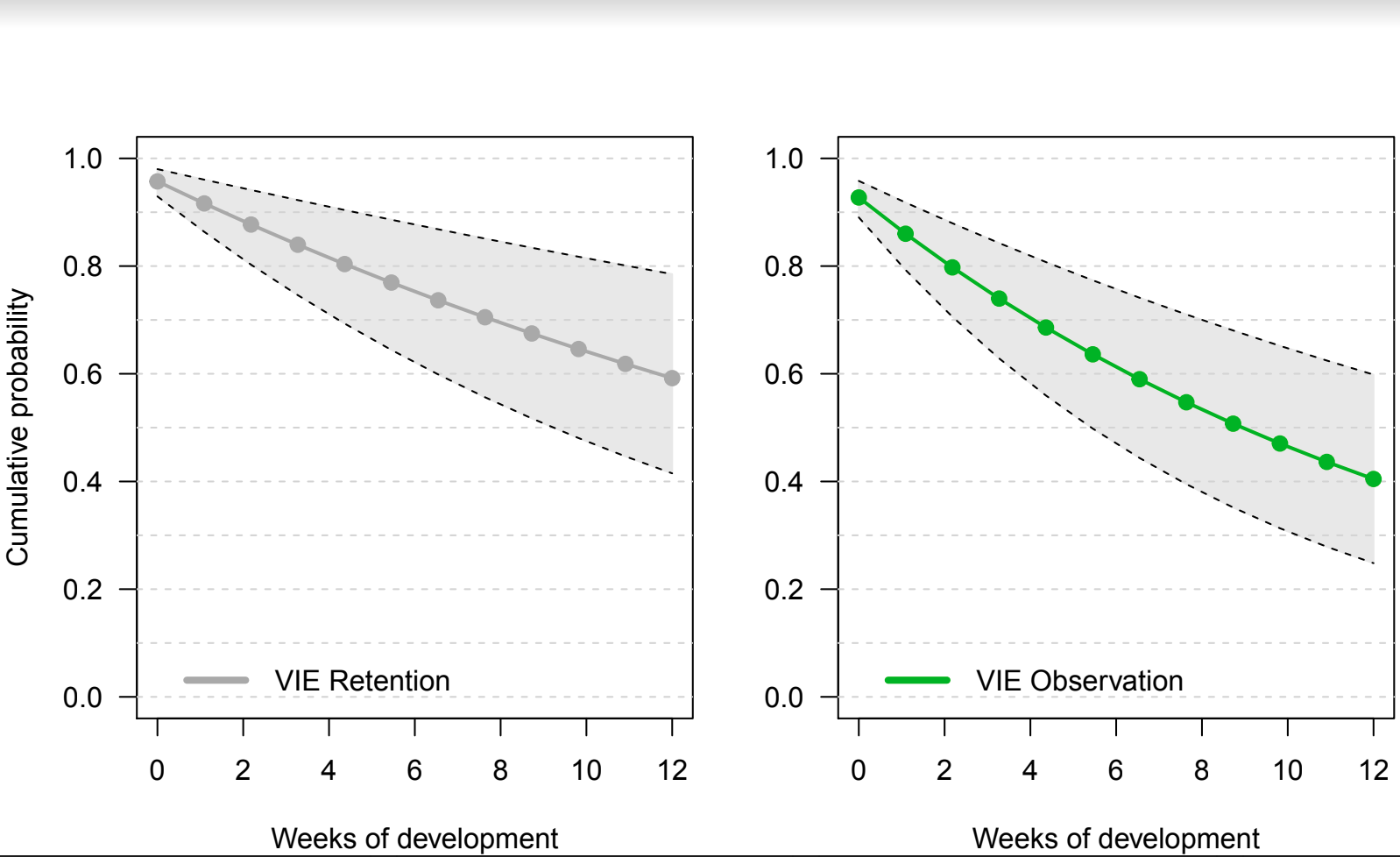

Supplementary figure 1. Posterior distribution and chain convergence of M1. Model assumed constant probabilities of observation and retention. Panel A is a trace plot indicating the convergence of 4 chains based on MCMC sampling and panel B is the posterior distribution generated by model function. Panel C visualizes the cumulative probability of observation across development; *ret* and *obs* denote retention and observation, respectively.

**A**.
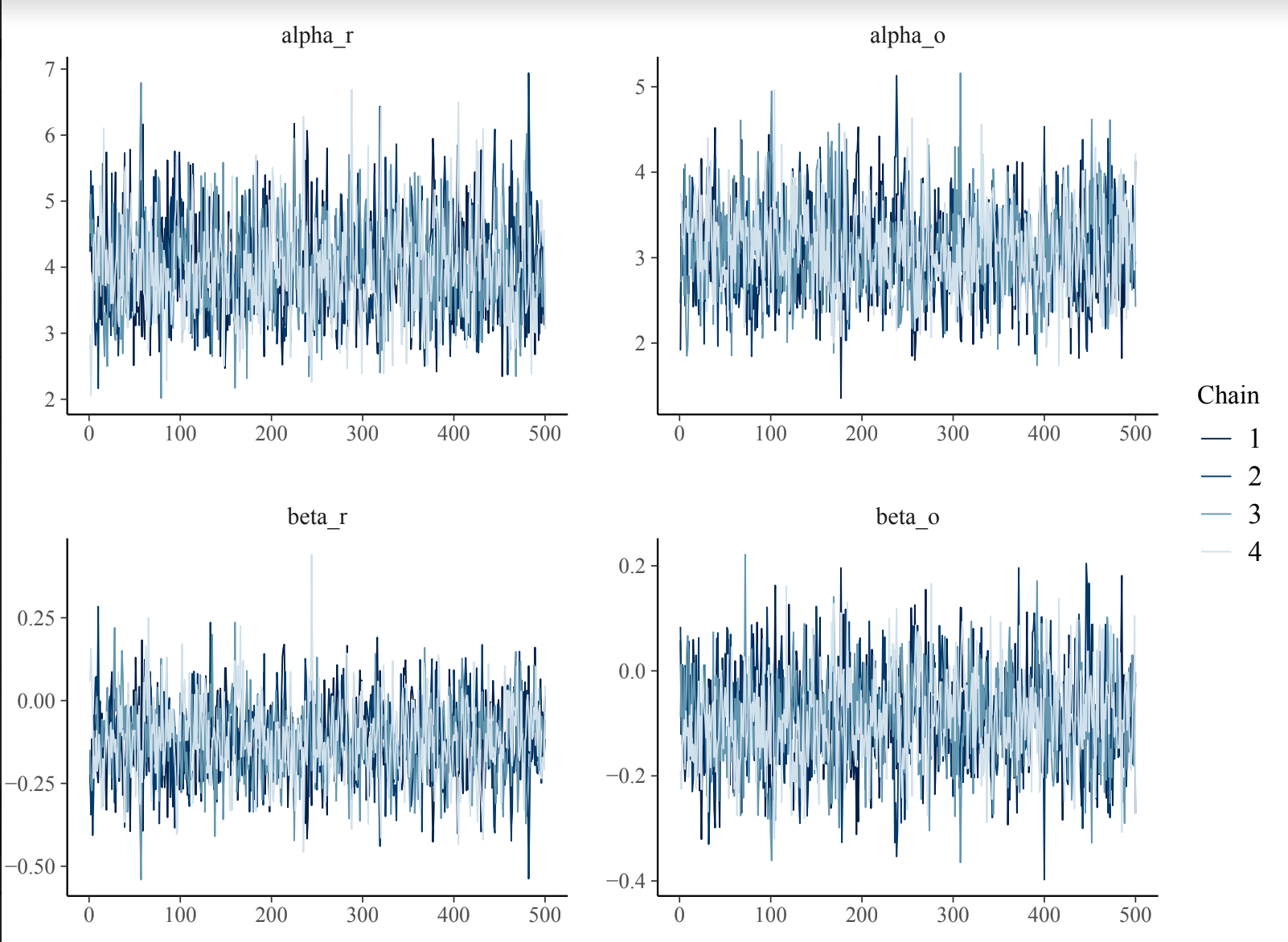
**B**.
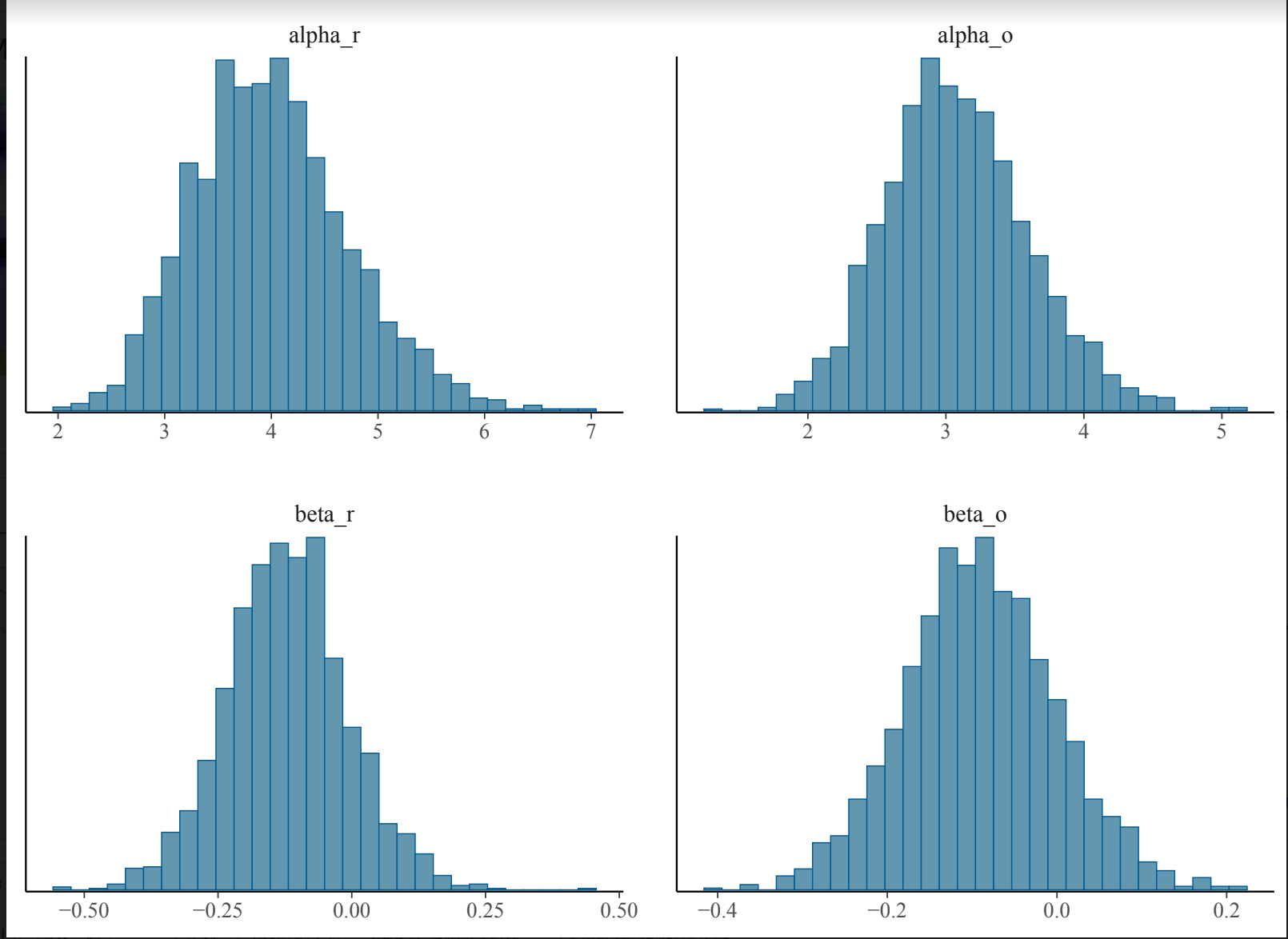

Supplementary figure 2. Posterior distribution and chain convergence of M2. Model considered a week effect on both probabilities using a logit link function. Panel A is a trace plot indicating the convergence of 4 chains based on MCMC sampling and panel B visualizes the posterior distribution generated by model function; *r* and *o* denote retention and observation, respectively.

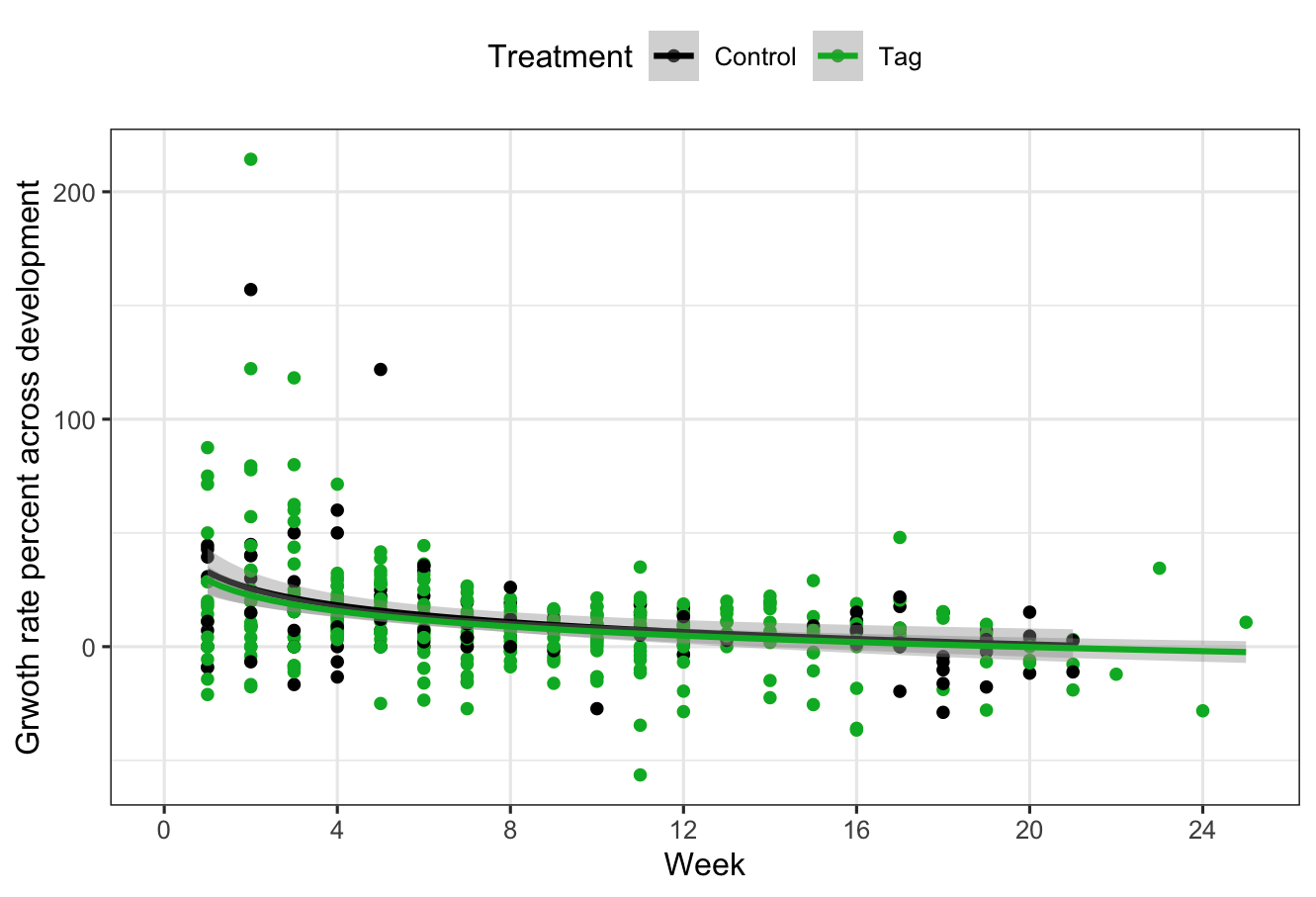

Supplementary figure 3. Weekly growth rate across development. Weekly growth rate decreases significantly over time. Grey bands represent 95% confidence intervals drawn by GLM smoother with a y ~log(x) formula.

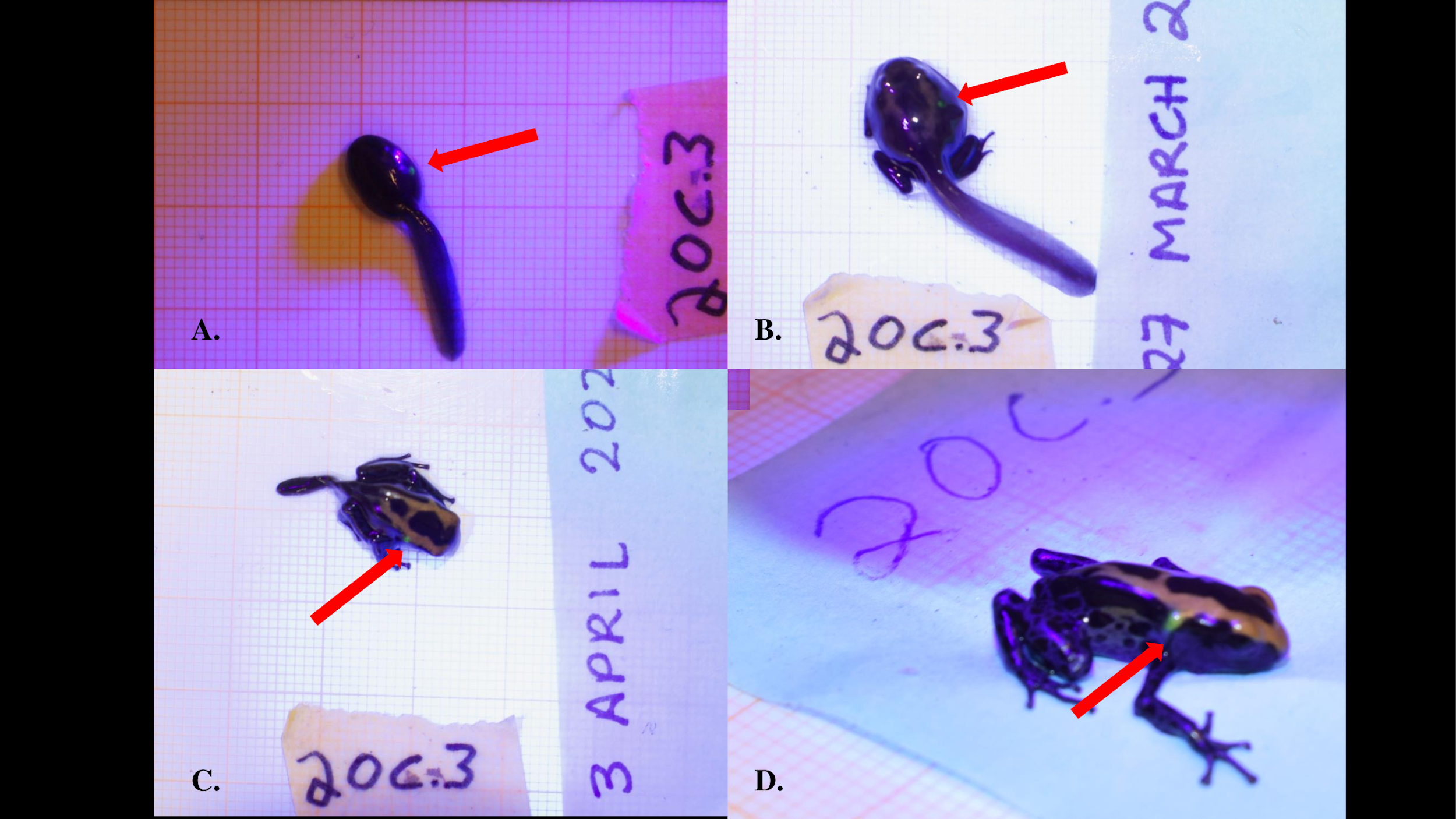

Supplementary figure 4. Fluorescent green VI Elastomer tag inserted dorsally on *Dendrobates tinctorius* shown on the same individual as (a) an early stage larvae, (b) a metamorph, (c) a late stage metamorph, and (d) a recently metamorphosed juvenile. All photos taken with Nikon DS5300 DSLR on 1 x 1 mm background under UV light to enhance tag detection.

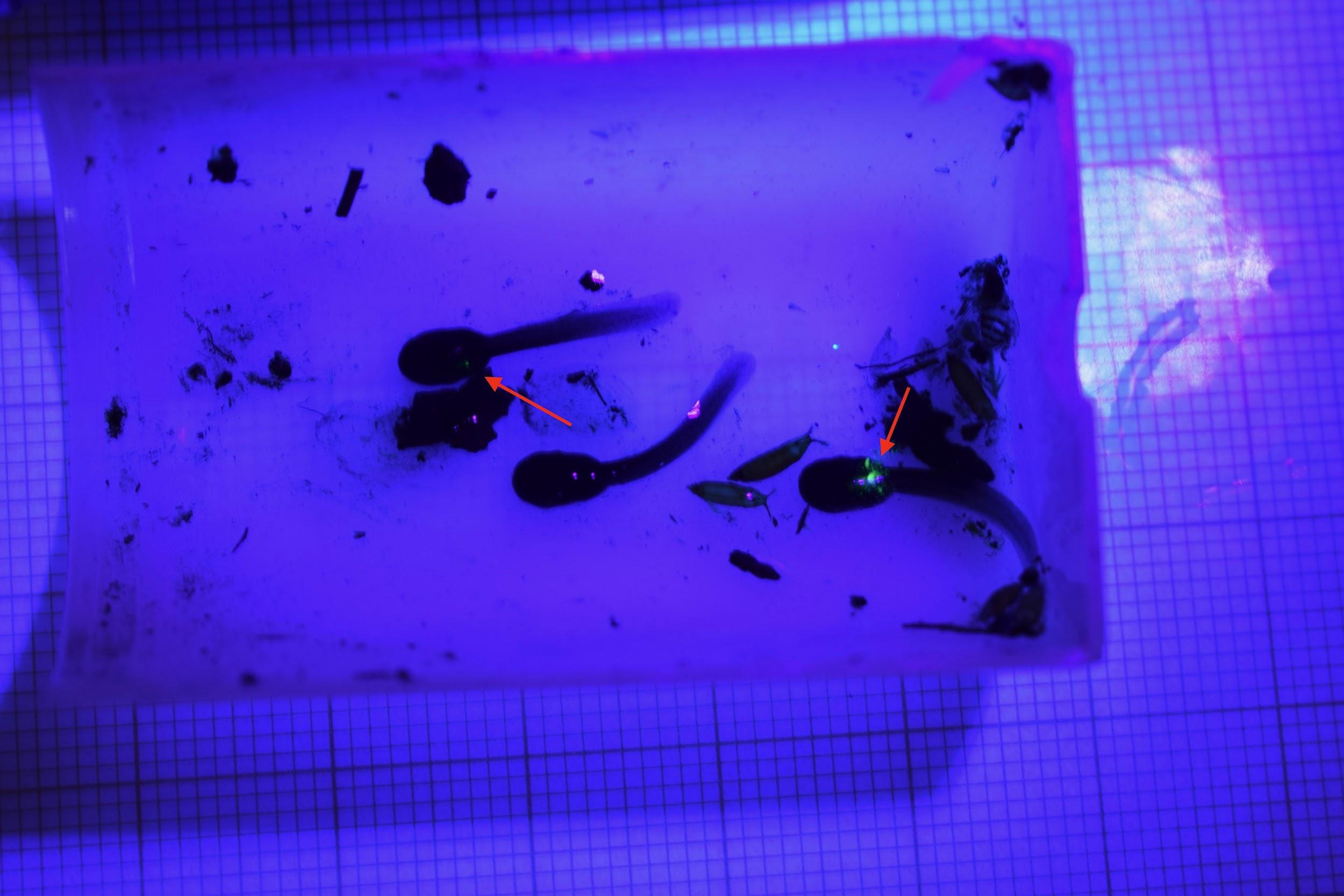

Supplementary figure 5. Recently hatched early term tadpoles that were successfully transported by father. Red arrows indicate tagged individuals.
