## Supplementary material for "Visible implant elastomer (VIE) success in early larval stages of a tropical amphibian species": VIE Growth/Survival Code

VIE_growth_survival.R

chloefouilloux

2020-04-28

```
###
## Elastomere Tagging Experiment ##
## Chloe Fouilloux, Guillermo Garcia, and Bibiana Rojas ##

#Code for Survival and Growth #


#first let's load some packages#
library(ggplot2)
library(dplyr)
##
## Attaching package: 'dplyr'
## The following objects are masked from 'package:stats':
##
##     filter, lag
## The following objects are masked from 'package:base':
##
##     intersect, setdiff, setequal, union
library(lme4)
## Loading required package: Matrix
library(survival)
library(survminer)
## Loading required package: ggpubr
## Loading required package: magrittr
library(lsmeans)
## Loading required package: emmeans
## The 'lsmeans' package is now basically a front end for 'emmeans'.
## Users are encouraged to switch the rest of the way.
## See help('transition') for more information, including how to
## convert old 'lsmeans' objects and scripts to work with 'emmeans'.
library(pbkrtest)
#Kenward Roger modification of F-tests for
#linear mixed effects models and a parametric bootstrap test
#for generalized linear mixed model
####
#Now, let's read in our most recent updated data file.
elastomere<- read.csv("Elastomer_Final.csv", header = T, sep = ",")

#############
###subsets###
#############
tagged <- elastomere[which(elastomere$Treatment=='Tag'), ] #Just tagged individuals
tagged_na <- tagged[!is.na(tagged$Tag_Y_N), ]   ##Tagged individuals with NAs dropped
threemonths<- elastomere[ which(elastomere$Week < 13), ] #subset larval time
tagged_na$Tag_0_1<- ifelse(tagged_na$Tag_Y_N == "Y", 1, 0)
##

############################################################
#########  Survival  #######################################
###########################################################
#### Below two graphs good at showing death across control and tagged groups
ggplot(elastomere, aes(x = Week, y = Weight, color = Dead))+
  geom_point()+
  facet_wrap(~Treatment)
## Warning: Removed 25 rows containing missing values (geom_point).
```

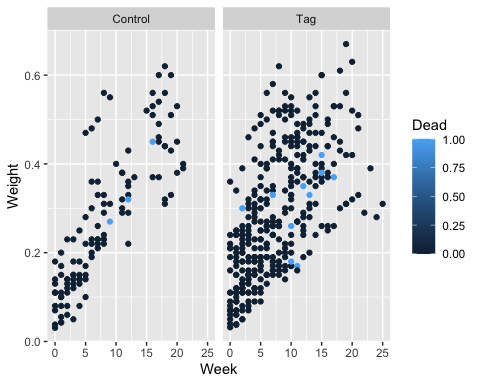

```
ggplot(elastomere, aes(x = Week, y = Weight, color = Dead))+
  geom_point(aes(shape = Treatment))+
  theme_bw()
## Warning: Removed 25 rows containing missing values (geom_point).
```

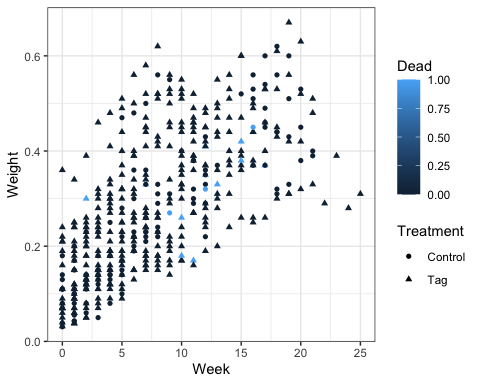

```
##Binomial Response Plot to Tag Survival Across all time
#Remember that this is skewed later in development because of all of our froglet death.
ggplot(elastomere, aes(x = Week, y = Dead, color = Treatment))+
  geom_point()+
  geom_smooth(method = "glm",
              method.args = list(family = "binomial"),aes(fill = Treatment),
              se = T, fullrange = T)+
  theme_bw()
## `geom_smooth()` using formula 'y ~ x'
```

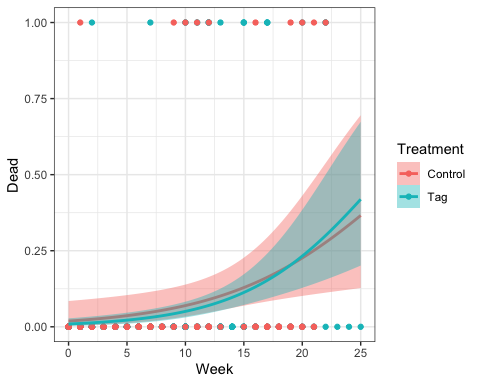

```
#If we subset this to just look at the first 3 months of development (larval)
##Binomial Response Plot to Tag Survival Across all time

#This plot is to look at all of the invidividuals tested, see whose dead and
#if I've entered any data incorrectly.
ggplot(threemonths, aes(x = Week, y = Dead, color = Treatment))+
  geom_point()+
  geom_smooth(method = "glm", formula=y~(x),
              method.args = list(family = "binomial"),aes(fill = Treatment),
              se = F, fullrange = T)+
  ylab("Larval death rate")+
  geom_hline(yintercept = 0.1, linetype = "dashed", alpha = 0.3)+
  theme_bw()+
  scale_x_continuous(breaks = seq(0, 12, 4))
```

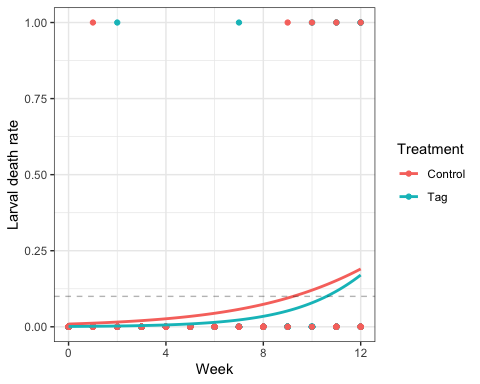

```
############################################################
#########  Calculating Growth Rate #########################
###########################################################

##Calculating average size at time of tagging
weight_cont<- elastomere[ which(elastomere$Week == 0 & elastomere$Treatment== "Control"), ]
weight_tag<- elastomere[ which(elastomere$Week == 0 & elastomere$Treatment== "Tag"), ]

mean(weight_cont$Weight, na.rm = T) ##0.099
## [1] 0.09989
min(weight_cont$Weight, na.rm = T) #0.0307
## [1] 0.0307
max(weight_cont$Weight, na.rm = T) #0.18
## [1] 0.18
mean(weight_tag$Weight, na.rm = T) ##0.12
## [1] 0.12559
min(weight_tag$Weight, na.rm = T) #0.0318
## [1] 0.0318
max(weight_tag$Weight, na.rm = T) #0.36
## [1] 0.36
#se
sd(weight_cont$Weight, na.rm=TRUE) /
  sqrt(length(weight_cont$Weight[!is.na(weight_cont$Weight)])) #0.0145
## [1] 0.01457609
sd(weight_tag$Weight, na.rm=TRUE) /
  sqrt(length(weight_tag$Weight[!is.na(weight_tag$Weight)])) #0.019
## [1] 0.0193278
#######################
### overall growth rate ###
#######################
growth_rate = elastomere %>%
  # first sort by week for each individual tested
  arrange(ID, Week) %>%
  group_by(ID) %>% #this is to make sure that the difference is just within an individualso that
  #last week of one does not become the first week of the next
  mutate(Diff_week = Week - lag(Week),  # Difference in time (just in case there are gaps)
         Diff_growth = Weight - lag(Weight), # Difference in weight between weeks
         Rate_percent = (Diff_growth / Diff_week)/lag(Weight) * 100) # growth rate in percent

#Visualizing growth rate#

#######################
#Growth rate percent across development#
###################
weekly<-
  ggplot(growth_rate, aes(color = Treatment, y = Rate_percent, x = Week))+
  geom_point()+
  stat_smooth(method = "glm", fullrange = F,
              formula = y ~ log(x))+
  ylab("Grwoth rate percent across development")+
  scale_x_continuous(breaks = seq(0, 24, 4))+
  theme_bw()+
  theme(legend.position = "top", panel.grid.minor.x = element_blank())


weekly+ scale_color_manual(values = c("black", "#0eb300"))
## Warning: Removed 64 rows containing non-finite values (stat_smooth).
## Warning: Removed 64 rows containing missing values (geom_point).
```

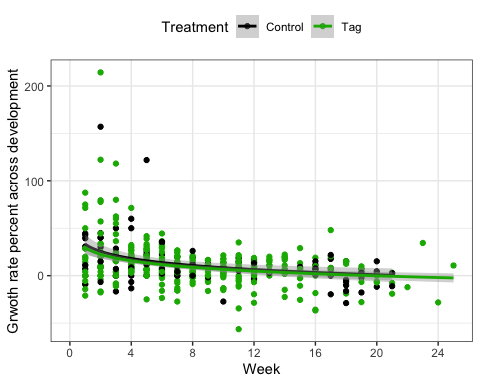

```
##############################################################
###### MODELLING  GROWTH RATE ###################################
#############################################################

#Looking at weight is great and all, but what we really want is growth RATE from
## week to week.
#not possion or gamma because of growth rate can be negative.
weight.sub<-lmer(Rate_percent ~ Treatment + (1|ID), data = growth_rate)
weight_week<- lmer(Rate_percent ~ Treatment* Week + (1|ID), data = growth_rate)
weightweek<- lmer(Rate_percent ~ Treatment+ Week + (1|ID), data = growth_rate) #best
weight.one.week<- lmer(Rate_percent ~ 1+ Week + (1|ID), data = growth_rate)
weight.one<-lmer(Rate_percent ~ 1 + (1|ID), data = growth_rate)
weight.lm<-lm(Rate_percent ~ Treatment, data = growth_rate)
weight.glm<-glm(Rate_percent ~ Treatment, data = growth_rate)

#AIC Model Comparison Showdown
aic<-AIC(weight.lm,
         weight.sub,
         weight.one,
         weight_week,
         weightweek,
         weight.one.week,
         weight.glm)
aic
##                 df      AIC
## weight.lm        3 3981.163
## weight.sub       4 3975.814
## weight.one       3 3978.177
## weight_week      6 3927.392
## weightweek       5 3925.560
## weight.one.week  4 3928.759
## weight.glm       3 3981.163
aic[order(aic$AIC),]
##                 df      AIC
## weightweek       5 3925.560
## weight_week      6 3927.392
## weight.one.week  4 3928.759
## weight.sub       4 3975.814
## weight.one       3 3978.177
## weight.lm        3 3981.163
## weight.glm       3 3981.163
##In this case, I would say that a delta(AIC) of 1.8 provides possible
##support for a model with a week*treatment interaction
##so let's explore both the top two models (weightweek) and (weight_week)

summary(weight_week) #interaction between weight and week
## Linear mixed model fit by REML ['lmerMod']
## Formula: Rate_percent ~ Treatment * Week + (1 | ID)
##    Data: growth_rate
##
## REML criterion at convergence: 3915.4
##
## Scaled residuals:
##     Min      1Q  Median      3Q     Max
## -2.8637 -0.5135 -0.0770  0.3627  8.7288
##
## Random effects:
##  Groups   Name        Variance Std.Dev.
##  ID       (Intercept)  23.01    4.797
##  Residual             476.94   21.839
## Number of obs: 434, groups:  ID, 38
##
## Fixed effects:
##                   Estimate Std. Error t value
## (Intercept)       26.70010    4.13731   6.453
## TreatmentTag      -3.76330    4.80092  -0.784
## Week              -1.54314    0.35776  -4.313
## TreatmentTag:Week  0.06013    0.43000   0.140
##
## Correlation of Fixed Effects:
##             (Intr) TrtmnT Week
## TreatmentTg -0.862
## Week        -0.771  0.664
## TrtmntTg:Wk  0.641 -0.771 -0.832
#Here we can see that the interaction term is not significant, so it's
#logical that we drop it from the model. Let's take a look at the next model!

summary(weightweek) #weight and week as addative effects
## Linear mixed model fit by REML ['lmerMod']
## Formula: Rate_percent ~ Treatment + Week + (1 | ID)
##    Data: growth_rate
##
## REML criterion at convergence: 3915.6
##
## Scaled residuals:
##     Min      1Q  Median      3Q     Max
## -2.8639 -0.5170 -0.0799  0.3660  8.7343
##
## Random effects:
##  Groups   Name        Variance Std.Dev.
##  ID       (Intercept)  23.17    4.813
##  Residual             475.72   21.811
## Number of obs: 434, groups:  ID, 38
##
## Fixed effects:
##              Estimate Std. Error t value
## (Intercept)   26.3323     3.1739   8.296
## TreatmentTag  -3.2484     3.0587  -1.062
## Week          -1.5019     0.1983  -7.575
##
## Correlation of Fixed Effects:
##             (Intr) TrtmnT
## TreatmentTg -0.752
## Week        -0.557  0.065
##Here we see that tagged individuals have an intercept that is -3.24 lower than controls.
##and that across weeks, tadpoles grow more slowly (-1.5)
#Let's look at how this pans out in an anova

anova(weightweek, ddf="Kenward-Roger")
## Analysis of Variance Table
##           Df  Sum Sq Mean Sq F value
## Treatment  1   156.2   156.2  0.3284
## Week       1 27295.1 27295.1 57.3759
################

## REPORTING LMM MODEL OUTPUT
##Show there is no difference in growth between the two treatments.
####################################
#### PLOTTING CONFIDENCE LIMITS OF GROWTH RATE#####
#################################
lsm<-lsmeans(weightweek, "Treatment") ##here we go!
lsm
##  Treatment lsmean   SE   df lower.CL upper.CL
##  Control     13.7 2.65 40.9     8.39     19.1
##  Tag         10.5 1.55 30.8     7.34     13.6
##
## Degrees-of-freedom method: kenward-roger
## Confidence level used: 0.95
with(summary(lsmeans(weightweek,spec="Treatment")),cbind(lower.CL,upper.CL))
##      lower.CL upper.CL
## [1,] 8.392251 19.07985
## [2,] 7.335148 13.64019
plot(lsm)
```

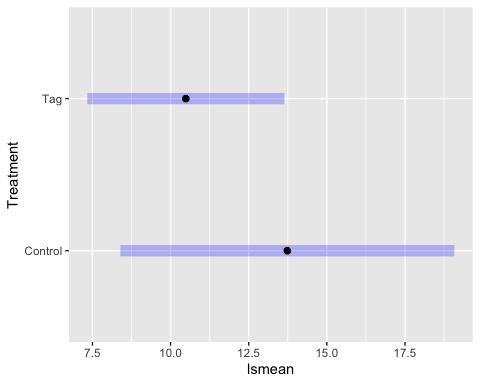

```
marginal = lsmeans(lsm,~ Treatment)

CLD<- multcomp::cld(marginal,
                    alpha=0.05,
                    Letters=letters,
                    adjust="tukey")
CLD
##  Treatment lsmean   SE   df lower.CL upper.CL .group
##  Tag         10.5 1.55 30.8     6.86     14.1  a
##  Control     13.7 2.65 40.9     7.59     19.9  a
##
## Degrees-of-freedom method: kenward-roger
## Confidence level used: 0.95
## Conf-level adjustment: sidak method for 2 estimates
## significance level used: alpha = 0.05
cld<-ggplot(CLD,
            aes(x= Treatment,
                y= lsmean,
                color = Treatment)) +
  geom_point(shape = 9,
             size = 4,
             position = position_dodge(0.4))+
  geom_errorbar(aes(ymin  =  lower.CL,
                    ymax  =  upper.CL),
                width =  0.2,
                size  =  0.7,
                position = position_dodge(0.4))+
  geom_text( aes( label = .group),
             nudge_y = 1.5,
             nudge_x = 0.05,
             fontface = "bold",
             size = 5.5)+
  ylab("Growth rate percent across development")+
  xlab("")+
  theme_bw()+
  theme(legend.position= "none",
        axis.text=element_text(size=12),
        axis.title.y = element_text(size = 12))

cld+ scale_color_manual(values = c("black", "#0eb300"))
```

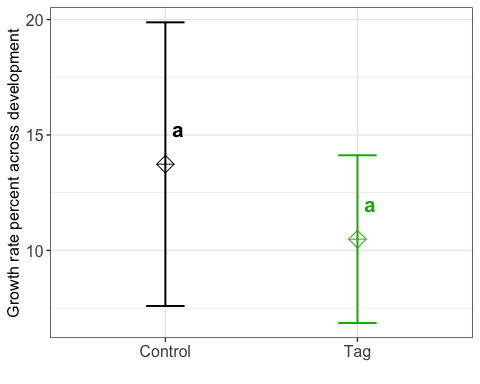

```
#CLs between both treatments overlap, which indicates that there is no difference
#between groups.


##############################################################
###### TAG EFFECT ON SURVIVAL ###################################
#############################################################

#Let's construct a survival curve!

alive <- with(elastomere, Surv(Week, Dead))
head(alive,80)
##  [1]  0+  1+  2+  3+  4+  5+  6+  7+  8+  9+ 10+ 13+ 14+ 15+ 16+ 17+ 18+ 19+ 20+
## [20] 21+ 22+ 23+ 24+ 25+  0+  1+  2+  3+  4+  5+  6+  7+  8+  9+ 10+ 13+ 14+ 15+
## [39] 16+ 17+ 18+ 19+ 20   0+  1+  2+  3+  4+  5+  6+  7+  9+ 10+ 11+ 12+ 13+ 14+
## [58] 15   0+  1+  2+  3+  4+  5+  6+  7+ 10+ 11+ 12+ 13+ 14+ 15+ 16+ 17   0+  1+
## [77]  2+  3+  4+  5+
alive_fit <- survfit(Surv(Week, Dead) ~ 1, data=elastomere)
summary(alive_fit, times = c(0, 4, 8, 12))
## Call: survfit(formula = Surv(Week, Dead) ~ 1, data = elastomere)
##
##  time n.risk n.event survival std.err lower 95% CI upper 95% CI
##     0    498       0    1.000  0.0000        1.000        1.000
##     4    353       2    0.995  0.0032        0.989        1.000
##     8    225       1    0.992  0.0050        0.982        1.000
##    12    132      12    0.916  0.0218        0.874        0.959
alive_trt_fit <- survfit(Surv(Week, Dead) ~ Treatment, data=elastomere)
summary(alive_trt_fit)
## Call: survfit(formula = Surv(Week, Dead) ~ Treatment, data = elastomere)
##
##                 Treatment=Control
##  time n.risk n.event survival std.err lower 95% CI upper 95% CI
##     1    114       1    0.991 0.00873        0.974        1.000
##     9     51       1    0.972 0.02106        0.931        1.000
##    10     48       1    0.952 0.02875        0.897        1.000
##    11     46       1    0.931 0.03478        0.865        1.000
##    12     40       1    0.908 0.04096        0.831        0.992
##    16     31       1    0.878 0.04900        0.787        0.980
##    19     14       1    0.816 0.07566        0.680        0.978
##    20      8       1    0.714 0.11609        0.519        0.982
##    21      4       1    0.535 0.17735        0.280        1.000
##    22      1       1    0.000     NaN           NA           NA
##
##                 Treatment=Tag
##  time n.risk n.event survival std.err lower 95% CI upper 95% CI
##     2    319       1    0.997 0.00313        0.991        1.000
##     7    192       1    0.992 0.00604        0.980        1.000
##    10    133       2    0.977 0.01204        0.953        1.000
##    11    110       2    0.959 0.01716        0.926        0.993
##    12     92       4    0.917 0.02618        0.867        0.970
##    13     74       1    0.905 0.02861        0.851        0.963
##    15     51       3    0.852 0.04017        0.776        0.934
##    17     32       3    0.772 0.05702        0.668        0.892
##    20     14       1    0.717 0.07501        0.584        0.880
##    22      6       2    0.478 0.14671        0.262        0.872
## So, here we are seeing the effects of our tadpoles dying later in life
## because of our poor care protocol-- though everyone is dying, they
## are dying equally across treartments, meaning that our tag isn't the thing thats'
## causing them to die. SO thats. . . better than nothing?

## Now we can also just look at this for larval developement,
## which is the threemonths data set that I had subsetted earlier

alive_fit_larval <- survfit(Surv(Week, Dead) ~ Treatment, data=threemonths)

#Great! Now let's get graphing kids!
ggsurvplot(alive_fit_larval,
           data = threemonths,
           palette = c("grey", "#0eb300"),
           linetype = "strata",
           conf.int = T,
           size = 1,
           legend.title = (""),
           legend.labs = c(" Control", "Tag"),
           legend = "top",
           font.legend = c(12),
           font.x = c(13),
           font.y = c(13),
           xlab= ("Time (weeks)"),
           xlim= c(0,12),
           ylim = c(50, 100),
           break.time.by = 4,
           fun = "pct",
           ggtheme= theme(panel.grid.minor.x = element_blank(),
                          panel.grid = element_line(colour = "#f3f3f2"),
                          panel.background = element_rect(fill = "white",colour = "black"),
                          axis.text=element_text(size=13)))
```

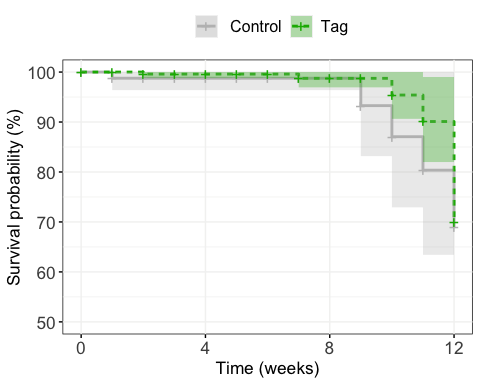

```
##############################################
## Fitting a Cox Proportional Hazards Model##
##############################################
cox <- coxph(Surv(Week, Dead) ~ Treatment, data = elastomere)
cox
## Call:
## coxph(formula = Surv(Week, Dead) ~ Treatment, data = elastomere)
##
##                 coef exp(coef) se(coef)      z     p
## TreatmentTag -0.1181    0.8886   0.3961 -0.298 0.766
##
## Likelihood ratio test=0.09  on 1 df, p=0.7669
## n= 498, number of events= 30
summary(cox)
## Call:
## coxph(formula = Surv(Week, Dead) ~ Treatment, data = elastomere)
##
##   n= 498, number of events= 30
##
##                 coef exp(coef) se(coef)      z Pr(>|z|)
## TreatmentTag -0.1181    0.8886   0.3961 -0.298    0.766
##
##              exp(coef) exp(-coef) lower .95 upper .95
## TreatmentTag    0.8886      1.125    0.4088     1.931
##
## Concordance= 0.525  (se = 0.063 )
## Likelihood ratio test= 0.09  on 1 df,   p=0.8
## Wald test            = 0.09  on 1 df,   p=0.8
## Score (logrank) test = 0.09  on 1 df,   p=0.8
anova(cox)
## Analysis of Deviance Table
##  Cox model: response is Surv(Week, Dead)
## Terms added sequentially (first to last)
##
##            loglik  Chisq Df Pr(>|Chi|)
## NULL      -129.91
## Treatment -129.86 0.0879  1     0.7669
cox1 <- coxph(Surv(Dead) ~ Treatment, data = elastomere)
summary(cox1)
## Call:
## coxph(formula = Surv(Dead) ~ Treatment, data = elastomere)
##
##   n= 498, number of events= 498
##
##                 coef exp(coef) se(coef)     z Pr(>|z|)
## TreatmentTag 0.07715   1.08021  0.10347 0.746    0.456
##
##              exp(coef) exp(-coef) lower .95 upper .95
## TreatmentTag      1.08     0.9257    0.8819     1.323
##
## Concordance= 0.544  (se = 0.044 )
## Likelihood ratio test= 0.56  on 1 df,   p=0.5
## Wald test            = 0.56  on 1 df,   p=0.5
## Score (logrank) test = 0.56  on 1 df,   p=0.5
anova(cox1)
## Analysis of Deviance Table
##  Cox model: response is Surv(Dead)
## Terms added sequentially (first to last)
##
##            loglik Chisq Df Pr(>|Chi|)
## NULL      -2598.9
## Treatment -2598.6 0.563  1      0.453
```
