## Supplementary material for "Visible implant elastomer (VIE) success in early larval stages of a tropical amphibian species": VIE Model Code

```
VIE Retention and Observation Experiment

Chloe Fouilloux, Guillermo Garcia-Costoya, Bibiana Rojas

#### BAYESIAN MODELS

# Load necessary packages
library(tidyverse)
library(reshape2)
library(R2jags)
library(bayesplot)
library(MCMCvis)
library(coda)

### PART 1: PREPARATION OF THE DATA

# Read the data
frog <- read.csv("Elastomer_Final.csv", header = T, sep = ",")

# Keep only individuals from "Tag" treatment
frog$treatment <- frog$Treatment
frog <- frog[which(frog$treatment=='Tag'), ]

# Tagged individuals with NAs dropped
frog <- frog[!is.na(frog$Tag_Y_N), ]

# Subset larval time
frog <- frog[ which(frog$Week < 12), ]

# Extract data into a simpler dataset
data <- mutate(frog,week=Week,observed=Tag_Y_N,id = ID)%>%select(week,id,observed)

# Modify the observation variable to have 1 & 0 instead of Y/N
data$obs <- ifelse(data$obs == "Y",1,0)

# Generate the "ret" column depedent on the observation colum.
data <- data %>% group_by(id) %>%
  mutate(ret=ifelse(max(week[which(obs==1)])>=week,1,NA)) %>% ungroup()


### PART 2: BAYESIAN MODELS

## COMMON STUFF FOR ALL MODELS

# Change id from factor to unique number
data$id <- as.numeric(as.factor(data$id))

# Retention matrix
ret_matrix <- acast(data,week ~ id, value.var = "ret")

## Fill in 1 in NA gaps
for (i in seq_len(ncol(ret_matrix))) {
  max_week <- max(which(ret_matrix[, i] == 1))
  ret_matrix[1:max_week, i] <- 1
}

# Observation matrix
obs_matrix  <- acast(data,week ~ id, value.var="obs")

# Model data in jags format
model_data <- list(n_inds = ncol(ret_matrix),n_weeks = nrow(ret_matrix),observation = obs_matrix)

# Options to run models
ni <- 1000000
nt <- 1000
nc <- 4


## MODEL WITH CONSTANT PROBABILITIES

# Model function
constant_prob_model <- function(){

  # Priors
  p_ret ~ dunif(0,1)
  p_obs ~ dunif(0,1)

  # Likelihood for the retention matrix (first week separated next weeks)
  for(i in 1:n_inds){
    retention[1,i]~ dbern(p_ret)
    for(w in 2:n_weeks){
      retention[w,i]~ dbern(retention[w-1,i]*p_ret)
    }
  }

  # Likelihood for the observation matrix
  for(i in 1:n_inds){
    for(w in 1:n_weeks){
      observation[w,i]~ dbern(retention[w,i]*p_obs)
    }
  }
}

# Model results run
constant_prob <- jags.parallel(data=model_data,model.file=constant_prob_model,parameters.to.save = c("p_ret","p_obs"),
                               inits = lapply(seq_len(1), function(i) {list(retention = ret_matrix)}),
                               n.chains =nc ,n.iter= 100000,n.thin= 10)

## MODEL WITH WEEK-DEPENDENT PROBABILITIES

# Model function
week_prob_model <- function(){

  # Prios
  alpha_r ~ dnorm(0,0.0001)
  beta_r ~ dnorm(0,0.0001)
  alpha_o ~ dnorm(0,0.0001)
  beta_o ~ dnorm(0,0.001)

  # Loop for the likelihood of the retention matrix
  for(i in 1:n_inds){

    # First week.
    p_ret[1, i] <- 1/(1 + exp(-alpha_r))
    retention[1,i] ~ dbern(p_ret[1, i])

    # Second week on, there is the beta and current state is dependant on previous state
    for(w in 2:n_weeks) {
      p_ret[w, i] <- 1/(1 + exp(-alpha_r - beta_r*(w-1)))
      retention[w,i] ~ dbern(retention[w-1,i]*p_ret[w, i])
    }
  }

  # Loop for the likelihood of the observation matrix
  for(i in 1:n_inds){
    for(w in 1:n_weeks){
      p_obs[w, i]<- 1/(1 + exp(-alpha_o - beta_o*(w-1)))
      observation[w,i] ~ dbern(retention[w,i]*p_obs[w, i])
    }
  }
}

# Model results run
week_prob <- jags.parallel(data=model_data,model.file=week_prob_model,parameters.to.save = c("alpha_r","beta_r","alpha_o","beta_o"),
                           init=lapply(seq_len(1),function(i){list(retention=ret_matrix)}),
                           n.chains = nc,n.iter = 100000,n.thin = 10)

## MODEL WITH WEEK DEPENDENT PROBABILITIES AND RANDOM EFFECT OF ID

# Model function
random_id_model <- function(){

  # General Priors
  alpha_ret ~ dnorm(0,0.0001)
  beta_ret ~ dnorm(0,0.0001)
  alpha_obs ~ dnorm(0,0.0001)
  beta_obs ~ dnorm(0,0.0001)

  # Priors for random effects
  s_alpha_ret_id ~ dnorm(0,100); T(0,)
  tau_alpha_ret_id <- pow(s_alpha_ret_id,-2)
  s_beta_ret_id ~ dnorm(0,100); T(0,)
  tau_beta_ret_id <- pow(s_beta_ret_id,-2)
  s_alpha_obs_id ~ dnorm(0,100); T(0,)
  tau_alpha_obs_id <- pow(s_alpha_obs_id,-2)
  s_beta_obs_id ~ dnorm(0,100); T(0,)
  tau_beta_obs_id <- pow(s_beta_obs_id,-2)

  # Loop for the likelihood of the retention matrix
  for(i in 1:n_inds){

    # Parameters for random effects
    alpha_ret_id[i] ~ dnorm(0,tau_alpha_ret_id)
    beta_ret_id[i] ~ dnorm(0,tau_beta_ret_id)

    # First week (logit link function + likelihood of the value whithin the matrix)
    p_ret[1, i] <- 1/(1 + exp(-alpha_ret -alpha_ret_id[i]))
    retention[1,i] ~ dbern(p_ret[1, i])

    # Second week on (same structure)
    for(w in 2:n_weeks) {
      p_ret[w, i] <- 1/(1 + exp(-alpha_ret -alpha_ret_id[i] - (beta_ret + beta_ret_id[i])*(w-1)))
      retention[w,i] ~ dbern(retention[w-1,i]*p_ret[w, i])
    }
  }

  # Loop for the likelihood of the observation matrix
  for(i in 1:n_inds){

    # Parameters for random effects
    alpha_obs_id[i] ~ dnorm(0,tau_alpha_obs_id)
    beta_obs_id[i] ~ dnorm(0,tau_beta_obs_id)

    # Loop for each week (same structure as before)
    for(w in 1:n_weeks){
      p_obs[w, i]<- 1/(1 + exp(-alpha_obs -alpha_obs_id[i] - (beta_obs + beta_obs_id[i])*(w-1)))
      observation[w,i] ~ dbern(retention[w,i]*p_obs[w, i])
    }
  }
}

# Model results run
random_id <- jags.parallel(data=model_data,model.file=random_id_model,parameters.to.save = c("alpha_ret","beta_ret","alpha_obs","beta_obs",
                                                                                              "s_alpha_ret_id","s_beta_ret_id","s_alpha_obs_id","s_beta_obs_id"),
                           init = lapply(seq_len(1), function(i) {list(retention = ret_matrix)}),n.chains=nc,n.iter=10000,n.thin=10)


### PART 3 DIC SCORES COMPARISON AND MODEL SELECTION

# DIC Values for all models got into one vector
cp <- constant_prob$BUGSoutput$DIC
wp <- week_prob$BUGSoutput$DIC
ri <- random_id$BUGSoutput$DIC

# DIC comparison dataframe
dic <- c(cp,wp,ri)
model <- c("Constant probability","Week probability","Random id effect")
delta_dic <- c(0,wp-cp,ri-cp)
comparison <- data.frame(model,dic,delta_dic)

### PART 4 CHAIN CONVERGENCE AND SAMPLING INDEPENDENCE

## MODEL WITH WEEK-DEPENDENT PROBABILITIES

# Change output into mcmc.list object
wprob <- as.mcmc.list(as.mcmc(week_prob))

# Potential scale reduction factor (PSRF)
gelman.diag(wprob)

# Number of independent samples
effectiveSize(wprob)


# VIE Retention and Observation Experiment

# Chloe Fouilloux, Guillermo Garcia-Costoya, Bibiana Rojas

#### PLOTS FOR MODELLING RESULTS

### PART 1: CONSTANT PROBABILITIES MODEL

# Summarized mcmc results
cprob <- coda::as.mcmc(constant_prob)[, c("p_ret", "p_obs")]

# PLOT 1: POSTERIOR DISTRIBUTIONS
mcmc_hist(cprob)

# PLOT 2: MCMC CHAIN TRACE
mcmc_trace(cprob)

# PLOT 3: DISCRETE PROBABILITIES

# Prepare the grid over the week range to estime the curves
n_weeks <- nrow(obs_matrix)
week_grid <- seq(0,n_weeks,length.out = 12)

# Convert MCMC output into tibble
zcp <- as_tibble(do.call(rbind,cprob))

# Plot paring instructions
par(mfrow=c(1,2),mar=c(4,4,4,0.25))

# PLOT FOR RETENTION PROBABILITY

# Fit lines for every probability
zcp[["pred_p_ret"]]<- pmap(list(zcp[["p_ret"]]),function(pret)tibble(week=week_grid,p_ret =pret))
# Extract all the curve values and put them into a matrix
cp_pred_p_ret <- lapply(zcp[["pred_p_ret"]],function(x) x %>% pull(p_ret)) %>% do.call(cbind,.)
# calculate quantiles to create the evelopes on the plot
cpr95 <- t(apply(cp_pred_p_ret,1,quantile,probs=c(0.025,0.975)))
# Draw credible interval
col_envelope <- adjustcolor("lightgrey",alpha.f=0.5)
abline(h=c(seq(0,1,0.1)),lty=2,col="lightgrey")
plot(0, xlim = c(0, n_weeks), ylim = c(0.8, 1), type = "n", xlab = "Weeks of development",
     ylab = "Discrete Probability", las = 1)
abline(h=c(seq(0.8,1,0.05)),lty=2,col="lightgrey")
polygon(x = c(week_grid, rev(week_grid)), y = c(cpr95[, 1], rev(cpr95[, 2])),col = col_envelope, border = col_envelope)
lines(cpr95[, 1]~ week_grid,lty=2)
lines(cpr95[, 2]~ week_grid,lty=2)
# Mean prediction line to the plot
cp_table <- MCMCsummary(constant_prob)
cp_p_ret <- rep(cp_table[3,1],length(week_grid))
lines(cp_p_ret ~ week_grid,type="o",lwd=2,pch=19,col="grey")
# Add legend
legend(x=0,y=0.83, lty=1, lwd=4,col = "grey",bty="n",legend = c("VIE Retention"))

# PLOT FOR OBSERVATION PROBABILITY

# Fit lines for every probability
zcp[["pred_p_obs"]]<- pmap(list(zcp[["p_obs"]]),function(pobs)tibble(week=week_grid,p_obs=pobs))
# Extract all the curve values and put them into a matrix
cp_pred_p_obs <- lapply(zcp[["pred_p_obs"]],function(x) x %>% pull(p_obs)) %>% do.call(cbind,.)
# calculate quantiles to create the evelopes on the plot
cpo95 <- t(apply(cp_pred_p_obs,1,quantile,probs=c(0.025,0.975)))
# Draw credible interval
plot(0, xlim = c(0, n_weeks), ylim = c(0.8, 1), type = "n", xlab = "Weeks of development",
     ylab = "",ylabt="n", las = 1)
abline(h=c(seq(0.8,1,0.05)),lty=2,col="lightgrey")
polygon(x = c(week_grid, rev(week_grid)), y = c(cpo95[, 1], rev(cpo95[, 2])),col = col_envelope, border = col_envelope)
lines(cpo95[, 1]~ week_grid,lty=2)
lines(cpo95[, 2]~ week_grid,lty=2)
# Mean prediction line to the plot
cp_p_obs <- rep(cp_table[2,1],length(week_grid))
lines(cp_p_obs ~ week_grid,type="o",lwd=2,pch=19,col="#0eb300")
# Add legend
legend(x=0,y=0.83, lty=1, lwd=4,col = "#0eb300",bty="n",legend = c("VIE Observation"))

# PLOT 4: CUMULATIVE PROBABILITIES

# PROBABILITY OF RETENTION

# Probability of retention matrix
pret_mat <- cumprod(zcp[["pred_p_ret"]][[1]][["p_ret"]])
for(i in 2:nrow(zcp)){pret_mat <- cbind(pret_mat,cumprod(zcp[["pred_p_ret"]][[i]][["p_ret"]]))}
# 95% quantile
cpcr95 <- t(apply(pret_mat,1,quantile,probs=c(0.025,0.975)))
# Draw credible interval
plot(0, xlim = c(0, n_weeks), ylim = c(0, 1), type = "n", xlab = "Weeks of development",
     ylab = "Cumulative probability", las = 1)
abline(h=c(seq(0,1,0.1)),lty=2,col="lightgrey")
polygon(x = c(week_grid, rev(week_grid)), y = c(cpcr95[, 1], rev(cpcr95[, 2])),col = col_envelope, border = col_envelope)
lines(cpcr95[, 1]~ week_grid,lty=2)
lines(cpcr95[, 2]~ week_grid,lty=2)
# Mean prediction line to the plot
cp_p_ret_cum <- cumprod(cp_p_ret)
lines(cp_p_ret_cum ~ week_grid,type="o",lwd=2,pch=19,col="darkgrey")
# Add legend
legend(x=0,y=0.1, lty=1, lwd=4,col = "darkgrey",bty="n",legend = c("VIE Retention"))

# PROBABILITY OF OBSERVATION

# Probability of retention matrix
pobs_mat <- cumprod(zcp[["pred_p_obs"]][[1]][["p_obs"]])
for(i in 2:nrow(zcp)){pobs_mat <- cbind(pobs_mat,cumprod(zcp[["pred_p_obs"]][[i]][["p_obs"]]))}
# 95% quantile
cpco95 <- t(apply(pobs_mat,1,quantile,probs=c(0.025,0.975)))
# Draw credible interval
plot(0, xlim = c(0, n_weeks), ylim = c(0, 1), type = "n", xlab = "Weeks of development",ylab = "",ylabt="n", las = 1)
abline(h=c(seq(0,1,0.1)),lty=2,col="lightgrey")
polygon(x = c(week_grid, rev(week_grid)), y = c(cpco95[, 1], rev(cpco95[, 2])),col = col_envelope, border = col_envelope)
lines(cpco95[, 1]~ week_grid,lty=2)
lines(cpco95[, 2]~ week_grid,lty=2)
# Mean prediction line to the plot
cp_p_obs_cum <- cumprod(cp_p_obs)
lines(cp_p_obs_cum ~ week_grid,type="o",lwd=2,pch=19,col="#0eb300")
# Add legend
legend(x=0,y=0.1, lty=1, lwd=4,col = "#0eb300",bty="n",legend = c("VIE Observation"))

### PART 2: WEEK-DEEPENDENT PROBABILITIES MODEL

# Summarized mcmc results
wprob <- coda::as.mcmc(week_prob)[, c("alpha_r", "alpha_o","beta_r","beta_o")]

# PLOT 1: POSTERIOR DISTRIBUTIONS
mcmc_hist(wprob)

# PLOT 2: MCMC CHAIN TRACE
mcmc_trace(wprob)

# PLOT 3: DISCRETE PROBABILITIES

# Convert MCMC output into tibble
zwp <- as_tibble(do.call(rbind,wprob))

# PLOT FOR RETENTION PROBABILITY

# Predicted probabilities of retention per iteration per week
zwp[["pred_p_ret"]]<-pmap(list(zwp[["alpha_r"]],zwp[["beta_r"]]),function(a,b)tibble(week=week_grid,p_ret=1/(1+exp(-(a+b*week_grid)))))
# Extract all curve values and get them into a single matrix
wp_pred_p_ret <- lapply(zwp[["pred_p_ret"]],function(x)x%>%pull(p_ret))%>%do.call(cbind,.)
# 95% envelope
wpr95 <- t(apply(wp_pred_p_ret,1,quantile,probs=c(0.025,0.975)))
# Draw credible interval
plot(0, xlim = c(0, n_weeks), ylim = c(0.5, 1), type = "n", xlab = "Weeks of development",
     ylab = "Discrete Probability", las = 1)
abline(h=c(seq(0.5,1,0.05)),lty=2,col="lightgrey")
polygon(x = c(week_grid, rev(week_grid)), y = c(wpr95[, 1], rev(wpr95[, 2])),col = col_envelope, border = col_envelope)
lines(wpr95[, 1]~ week_grid,lty=2)
lines(wpr95[, 2]~ week_grid,lty=2)
# Mean prediction line to the plot
wp_table <- MCMCsummary(week_prob)
wp_p_ret <- 1/(1+exp(-wp_table[2,1]-(wp_table[4,1])*week_grid))
lines(wp_p_ret ~ week_grid,type="o",lwd=2,pch=19,col="darkgrey")
# Add legend
legend(x=0,y=0.55, lty=1, lwd=4,col = "darkgrey",bty="n",legend = c("VIE Retention"))


# PLOT FOR OBSERVATION PROBABILITY

# Predicted probabilities of retention per iteration per week
zwp[["pred_p_obs"]]<-pmap(list(zwp[["alpha_o"]],zwp[["beta_o"]]),function(a,b)tibble(week=week_grid,p_obs=1/(1+exp(-(a+b*week_grid)))))
# Extract all curve values and get them into a single matrix
wp_pred_p_obs <- lapply(zwp[["pred_p_obs"]],function(x)x%>%pull(p_obs))%>%do.call(cbind,.)
# 95% envelope
wpo95 <- t(apply(wp_pred_p_obs,1,quantile,probs=c(0.025,0.975)))
# Draw credible interval
plot(0, xlim = c(0, n_weeks), ylim = c(0.5, 1), type = "n", xlab = "Weeks of development",
     ylab = "",ylabt="n",las = 1)
abline(h=c(seq(0.5,1,0.05)),lty=2,col="lightgrey")
polygon(x = c(week_grid, rev(week_grid)), y = c(wpo95[, 1], rev(wpo95[, 2])),col = col_envelope, border = col_envelope)
lines(wpo95[, 1]~ week_grid,lty=2)
lines(wpo95[, 2]~ week_grid,lty=2)
# Mean prediction line to the plot
wp_table <- MCMCsummary(week_prob)
wp_p_obs <- 1/(1+exp(-wp_table[1,1]-(wp_table[3,1])*week_grid))
lines(wp_p_obs ~ week_grid,type="o",lwd=2,pch=19,col="#0eb300")
# Add legend
legend(x=0,y=0.55, lty=1, lwd=4,col = "#0eb300",bty="n",legend = c("VIE Observation"))

# PLOT 4: CUMULATIVE PROBABILITIES


# PLOT FOR PROBABILITY OF RETENTION

# PROBABILITY OF RETENTION

# Matrix of all cumulative proabbilities
pret_mat <- cumprod(zwp[["pred_p_ret"]][[1]][["p_ret"]])
for(i in 2:nrow(zwp)){pret_mat <- cbind(pret_mat,cumprod(zwp[["pred_p_ret"]][[i]][["p_ret"]]))}
# Quantile 95
wpcr95 <- t(apply(pret_mat,1,quantile,probs=c(0.025,0.975)))
# Drawing envelope
plot(0, xlim = c(0, n_weeks), ylim = c(0, 1), type = "n", xlab = "Weeks of development",
     ylab = "Cumulative probability", las = 1)
abline(h=c(seq(0,1,0.1)),xlim=c(0,max(week_grid)),lty=2,col="lightgrey")
polygon(x = c(week_grid, rev(week_grid)), y = c(wpcr95[, 1], rev(wpcr95[, 2])),
        col = col_envelope, border = "lightgrey")
lines(wpcr95[, 1]~ week_grid,lty=2)
lines(wpcr95[, 2]~ week_grid,lty=2)
# Add the mean line
wp_cum_p_ret <- cumprod(wp_p_ret)
lines(wp_cum_p_ret ~ week_grid,type="o",lwd=2,pch=19,col="darkgrey")
legend(x=0,y=0.1, lty=1, lwd=4,col = "darkgrey",bty="n",legend = c("VIE Retention"))

# PROBABILITY OF OBSERVATION

# Matrix of all cumulative proabbilities
pobs_mat <- cumprod(zwp[["pred_p_obs"]][[1]][["p_obs"]])
for(i in 2:nrow(zwp)){pobs_mat <- cbind(pobs_mat,cumprod(zwp[["pred_p_obs"]][[i]][["p_obs"]]))}
# Quantile 95
wpco95 <- t(apply(pobs_mat,1,quantile,probs=c(0.025,0.975)))
# Drawing envelope
plot(0, xlim = c(0, n_weeks), ylim = c(0, 1), type = "n", xlab = "Weeks of development",
     ylab = "Cumulative probability", las = 1)
abline(h=c(seq(0,1,0.1)),xlim=c(0,max(week_grid)),lty=2,col="lightgrey")
polygon(x = c(week_grid, rev(week_grid)), y = c(wpco95[, 1], rev(wpco95[, 2])),
        col = col_envelope, border = "lightgrey")
lines(wpco95[, 1]~ week_grid,lty=2)
lines(wpco95[, 2]~ week_grid,lty=2)
# Add the mean line
wp_cum_p_obs <- cumprod(wp_p_obs)
lines(wp_cum_p_obs ~ week_grid,type="o",lwd=2,pch=19,col="#0eb300")
legend(x=0,y=0.1, lty=1, lwd=4,col = "#0eb300",bty="n",legend = c("VIE Observation"))
```
